# Structural basis for the evolution of a domesticated group II intron-like reverse transcriptase to function in host cell DNA repair

**DOI:** 10.1101/2025.01.14.632616

**Authors:** Seung Kuk Park, Mo Guo, Jennifer L. Stamos, Wantae Kim, Sidae Lee, Y. Jessie Zhang, Alan M. Lambowitz

## Abstract

A previous study found that a domesticated bacterial group II intron-like reverse transcriptase (G2L4 RT) functions in double-strand break repair (DSBR) via microhomology-mediated end joining (MMEJ) and that a mobile group II intron-encoded RT has a basal DSBR activity that uses conserved structural features of non-LTR-retroelement RTs. Here, we determined G2L4 RT apoenzyme and snap-back DNA synthesis structures revealing novel structural adaptations that optimized its cellular function in DSBR. These included a unique RT3a structure that stabilizes the apoenzyme in an inactive conformation until encountering an appropriate substrate; a longer N-terminal extension/RT0-loop with conserved residues that together with a modified active site favors strand annealing; and a conserved dimer interface that localizes G2L4 RT homodimers to DSBR sites with both monomers positioned for MMEJ. Our findings reveal how a non-LTR-retroelement RT evolved a dedicated cellular function and suggest new ways of optimizing these RTs for genome engineering applications.

## Introduction

Reverse transcriptases (RTs) are ancient enzymes that evolved from an RNA-dependent RNA polymerase thereby enabling a transition from an RNA to a DNA world^1–4^. Present day RTs are prevalent in bacteria, both as RTs encoded by mobile group II introns and as chromosomally encoded RTs that evolved from group II intron RTs to perform cellular functions, a process referred to as domestication^5–8^. Mobile group II introns are hypothesized to have entered ancestral eukaryotes with bacterial endosymbionts that gave rise to mitochondria and chloroplasts, proliferated in what became the nuclear genome, and evolved into both the core of the eukaryotic splicing apparatus (snRNAs U2, U5 and U6 and spliceosomal protein Prp8) and other eukaryotic RTs^9–13^. The latter include non-LTR-retrotransposon RTs, a family of enzymes that includes human LINE-1 element RT and whose RT core remains closely related to that of group II intron-encoded RTs^14,15^, followed by more divergent telomerase and retroviral RTs^13^.

Genome sequencing revealed that bacteria harbor a wide variety of chromosomally encoded group II intron-like RTs that are no longer associated with group II intron RNAs and have diversified over billions of years to perform a variety of cellular functions^5–8^. These include phage defense by different mechanisms, host-phage tropism switching, site-specific integration of RNA protospacers into CRISPR arrays, DSBR via MMEJ, and phage defense by *de novo* synthesis of new genes using the proficient template-switching activity of group II intron-like RTs^16–23^. In many cases, these cellular functions use novel biochemical mechanisms that could not have been rationally designed or even imagined based on existing knowledge.

All RTs share a common structural framework comprised of fingers, palm, and thumb or thumb surrogate regions that fold into a hand-like structure, forming a cleft containing an RT active site comprised of three conserved aspartate residues that bind catalytic Mg^2+^ ions^1,24–26^. Two of these aspartates are part of a conserved Y/FxDD motif found in all RTs. The RT fingers and palm contain 7 conserved sequence blocks (RT1-7) that include or are positioned around the RT active site to contribute to RT activity (Figure 1A)^25,26^. Bacterial and other prokaryotic RTs, mitochondrial retroplasmid RTs, and eukaryotic non-LTR-retrotransposon RTs, collectively termed non-LTR-retroelement RTs, contain three additional conserved regions: an N-terminal extension (NTE) with an RT0 loop and two expanded regions (RT2a and RT3a) between conserved RT sequence blocks. In group II intron RTs, these regions contribute to tighter binding pockets that enable higher fidelity and processivity than retroviral RTs^3,4,27^. Structural cognates of the NTE/RT0 loop, RT2a and RT3a are also present and functionally important in RNA-dependent RNA polymerases (RdRPs), evolutionary ancestors of RTs, but have been lost from retroviral RTs, which evolved a looser architecture and more promiscuous lifestyle to introduce and propagate mutational variations that help retroviruses evade host defense (Figure 1A)^4,27^.

**Figure 1.**
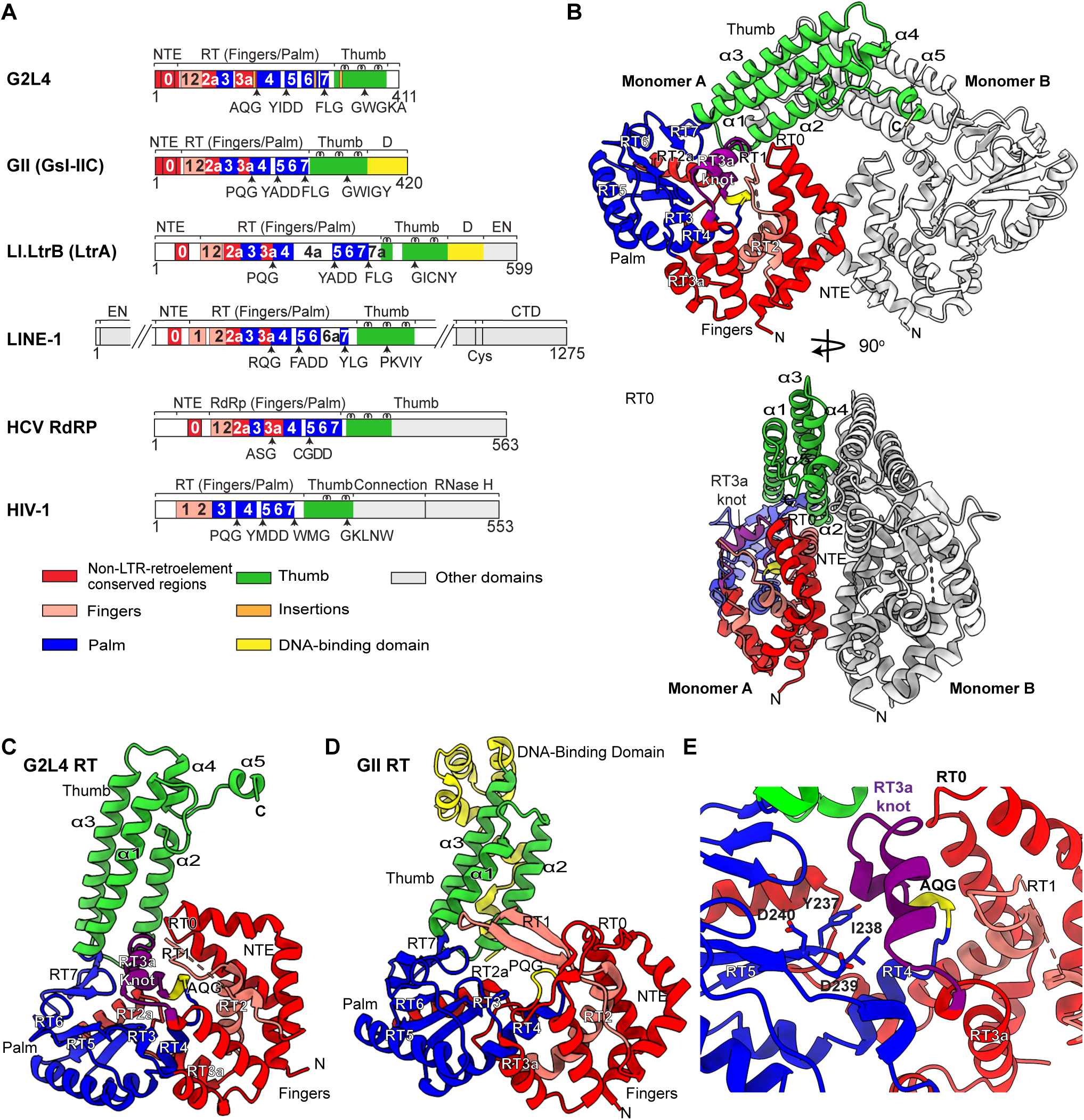
Structure of G2L4 RT apoenzyme. (A) Schematics showing the domain organization of G2L4 RT, Group IIC intron GsI-IIC (denoted GII) RT, Group IIA intron Ll.LtrB RT (LtrA protein), and human LINE-1 RT compared to Hepatitis C Virus (HCV) RdRP and retrovirus HIV-1 RT. RT1 to 7 are conserved sequence blocks in the fingers (salmon) and palm (blue) regions of all RTs. An N-terminal extension (NTE) with an RT0 loop, RT2a, and RT3a (red) are additional functionally important regions in non-LTR-retroelement RTs and RdRPs, but absent in retroviral RTs. (B) X-ray crystal structures of a G2L4 RT apoenzyme homodimer with a 90° rotating view. Monomer A regions are colored as in the schematic of panel A, and monomer B is colored gray. The AQG motif is highlighted in yellow, and the disordered RT1 is represented by a salmon-colored dashed line in monomer A. N indicates the N-terminus of each monomer. (C) Structure of the G2L4 RT apoenzyme monomer. (D) Structure of a monomeric GII RT in active conformation (PDB: 6AR1)^4^ in the same orientation as the G2L4 RT apoenzyme monomer in panel C. The bound template/primer and incoming dNTP in the structure are not shown. (E) G2L4 RT active-site region blocked by the RT3a knot in the apoenzyme structure. Active site YIDD residues are shown as sticks, and the RT3a knot is shown in purple.

Here, we focused on G2L4 RT, a bacterial group II intron-like RT that evolved to function in DSBR via MMEJ^20^, and determined a 2.6-Å crystal structure of full-length G2L4 RT apoenzyme and a 2.8-Å co-crystal structure of G2L4 RT bound in an active conformation to a snapback DNA synthesis substrate and incoming dNTP. The structures revealed a series of structural adaptations that evolved to optimize the cellular DSBR function of G2L4 RT, including I instead of A in the Y/FxDD motif at the RT active site and distinctive modifications in the NTE/RT0 loop, RT2a, and RT3a. Our findings enabled a detailed structural model of how G2L4 RT functions in DSBR and draw attention to these three distinctive regions as sites that can be modified to optimize non-LTR-retroelement RTs for biotechnological applications.

## Results

### G2L4 RT apoenzyme structure

Group II intron-encoded RTs typically bind group II intron RNAs co-transcriptionally to promote RNA splicing and tend to be unstable when removed from the intron RNA^28,29^. By contrast, domesticated group II intron-like RTs are synthesized as free-standing proteins that must bind their physiological substrates at different intracellular locations while avoiding deleterious interactions with cellular RNAs. To investigate how this might be done by G2L4 RT, we obtained a crystal structure of the full-length G2L4 RT apoenzyme (Figure 1B). The structure was solved at 2.60-Å resolution in space group P2_1_ by *de novo* phasing using selenomethionine. Unlike group II intron RTs, which typically appear in crystal or cryo-EM structures as monomers^4,30–36^, the G2L4 RT apoenzyme structure revealed two monomers per asymmetric unit forming a symmetric dimer, a difference verified by size-exclusion chromatography comparing G2L4 RT and a *Geobacillus stearothermopolis* group II intron RT (GsI-IIC RT denoted GII RT; Figure S1A). Continuous density could be traced for all amino acids, except those in two putative loops from Q65 to R78 in RT1 and from G116 to D120 in RT2a (Figure 1B and 1C).

The two G2L4 RT monomers that form the apoenzyme dimer have nearly identical hand-like folds comprised of fingers, palm, and thumb regions (Root Mean Square Deviation (RMSD) = 0.83 Å for main chain carbon atoms; Figure S1B), but lack DNA-binding (D) or DNA endonuclease (En) domains found in group II intron RTs (Figures 1A-D). Although G2L4 RT has relatively low overall sequence identity compared to GII RT (EMBOSS Needle sequence identity = 21.5%; Figure S1C)^37^, its tertiary structure is similar to that of GII RT (Figures 1C and 1D).

As in GII and other non-LTR-retroelement RTs, the fingers and palm of G2L4 RT contain 7 conserved RT sequence blocks (RT1-7) plus an NTE with an RT0 loop and extended regions RT2a and RT3a between the conserved RT sequence blocks (Figures 1A and S1C). The thumb domain of G2L4 RT is similar to that of GII RT in being comprised of 3 parallel α-helices (α1-3), but differs in extending straight up for 41 Å, 15 Å farther than in GII RT, and ending with additional short α-helices (α4 and α5) that interact with the thumb domain of the opposite monomer (Figures 1B-D)^4^.

The structures of most of the conserved RT motifs and the active site of G2L4 RT apoenzyme are similar to those of GII RT in an active conformation with bound substrates^4^. Two exceptions were: (i) RT1, which ordinarily forms two antiparallel β-strands over the active site, but was disordered and thus not visible in the G2L4 apoenzyme structure (represented by a salmon dashed line in Figure 1C), and (ii) the XQG motif (AQG in G2L4, PQG in GII RT), a critical element of the RT active site that functions in dNTP binding^4^, but is folded inward in the G2L4 RT apoenzyme structure compared to the active conformation of GII RT (Figures 1C and 1D).

As in GII RT, the NTE of the G2L4 RT apoenzyme is composed of two bent α-helices separated by a sharp turn that forms an RT0 loop, but with the RT0 loop smaller and the bent α-helices longer and closer to the RT active site than in GII RT (Figures 1C, 1D and S1C and see below). RT2a is nearly identical to that in the GII RT co-crystal structure, but with a hinge region (G116 to D120) that connects to RT2 and is disordered in the absence of bound substrate (Figures 1C and 1D). By contrast, RT3a in G2L4 RT differs markedly from that in group II intron RTs in forming a novel structural feature, a double-helical knot (denoted RT3a knot; highlighted in purple in Figures 1C-1E). The RT3a knot is a structural adaptation that completely blocks the active site of G2L4 RT, rendering the apoenzyme totally inactive until it encounters a physiological substrate, as described further below.

### Interaction of the RT3a knot with the G2L4 RT active site

The RT3a knot is comprised of 21 amino acids (aa) at the distal end of RT3a extending from N185 up to V205 just before the conserved AQG of the RT4 motif, and it includes a 9-aa insertion that is conserved in G2L4 RTs but absent in group II intron or other non-LTR-retroelement RTs (Figures 2A and S2A). These 21 amino acids form two α-helices that plug into the RT active site flush against the conserved YIDD at the catalytic center (Figures 1E and 2B), replacing a bound catalytic Mg^2+^ ion seen at this location in the active structures of GII and other RTs^4,15,38^. The RT3a knot was not predicted by any folding prediction program, including the most recent version of AlphaFold (AlphaFold 3.0, https://alphafold.ebi.ac.uk/)^39^, indicating that it was beyond the imagination of artificial intelligence. A strong interaction of the RT3a knot with the G2L4 RT active site is likely what displaced the RT1 β-strands that normally occupy this region causing them to become disordered and pulled the adjacent AQG motif inward where it became inaccessible to bulk solvent, as judged by modeled accessibility of a hypothetical water molecule.

**Figure 2.**
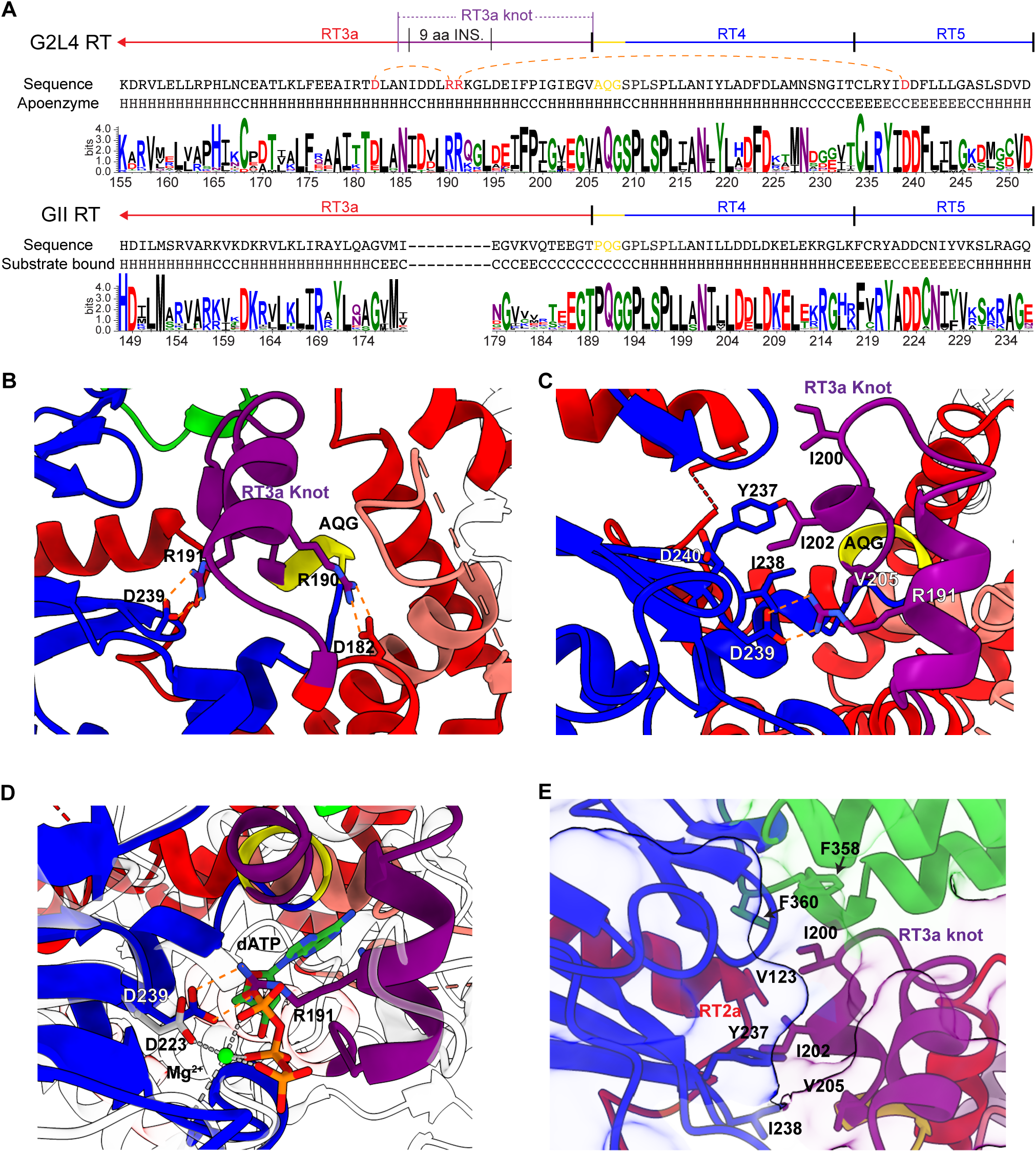
Amino acid sequence conservation and structure of the G2L4 RT RT3a knot. (A) WebLogos of the C-terminal portion of RT3a containing the RT3a knot, RT4, and RT5 of G2L4 RTs and the corresponding region of GII RTs. WebLogos are based on 130 G2L4 RTs and 500 GsI-IIC RTs identified by BLASTP^47^ as having ≥50% amino acid sequence identity aligned by ClustalW. Interacting residues described in the text are highlighted in red and connected by orange dashed lines. Amino acids in helical (H), random coil (C), and b-sheet (E) regions in the apoenzyme crystal structure are shown below the amino acid sequences. (B) Salt bridges between conserved residues in the RT3a knot and active site of G2L4 RT apoenzyme. Interacting residues arginine 190 and 191 in the RT3a knot and aspartate 182 and 239 in RT3a and RT5, respectively, are shown as sticks with salt bridges depicted as orange dashed lines. (C) Interactions of the RT3a knot (purple) with YIDD at the G2L4 RT active site. Key residues and salt bridges are depicted as in panel B. (D) Superimposition of active-site region of G2L4 RT apoenzyme (color-coded as in Figure 1A) and GII RT (gray/white) in an active conformation with a catalytic Mg^2+^ (green sphere) that interacts with GII RT D223 and an incoming dATP (dashed gray lines). Salt bridges of the corresponding aspartate (D239) with R191 in the RT3a knot of G2L4 RT apoenzyme structure are shown as orange dashed lines. (E) Hydrophobic interactions between the RT3a knot and the active site and thumb domain of G2L4 RT. Side chains of hydrophobic residues at the interface are shown as sticks. The surface of protein is represented by lightly shaded surface overlays demarcated by thin colored lines.

The closest resemblance to the interaction the RT3a of G2L4 RT with the active site was seen in apoenzyme structures of *Roseburia intestinalis* and *Eubacterium rectale* group II intron RT fingers and palm regions obtained with crystallographic constructs lacking a thumb domain (PDB code: 5HHJ and 5HHL; Figures S2A-S2C)^31^. In both of those structures, the distal end of the shorter RT3a, which lacks the 9-aa insertion, forms a one and a half turn α-helix that partially covers the active site YADD motif, but in an orientation that differs from that in G2L4 RT and does not extend to displace the RT1 β-strands, which remain well-ordered over the active-site pocket (Figures 2C and S2C)^31^. Thus, the 9-aa insertion is a unique evolutionary adaptation that enables complete blockage of the active site by RT3a in G2L4 RT.

The structure of the RT3a knot and its binding in the active-site pocket of G2L4 RT are stabilized by an extensive interaction network that includes salt bridges formed by two conserved arginine residues (R190 and R191) that are located within the 9-aa insertion and form part of the first α-helix of the RT3a knot (Figures 2A-2C). The first salt bridge is between R190 of the RT3a insertion and D182 located upstream in RT3a (Figure 2B). Notably, D182 is a conserved acidic residue in RT3a of G2L4 RTs, but a conserved glycine (G175) in RT3a of GII RTs (Figure 2A), indicating a functionally important covariation that stabilizes the RT3a knot. The second salt bridge is between R191 of the RT3a insertion and the conserved D239 of the YIDD motif at the active site (Figures 2B and 2C). In addition to preventing binding of an incoming nucleotide, this salt bridge replaces the catalytic Mg^2+^ seen at the corresponding aspartate residue in the active site in the crystal structure of GII RT^4^ (Figure 2D), thereby rendering G2L4 RT inactive.

The interactions of the second α-helix of the RT3a knot with the G2L4 RT active site are largely hydrophobic (Figure 2E). These interactions involve a series of greasy residues (I200, I202, V205) on a face of the helix that packs along a hydrophobic tunnel formed by Y237 and I238 of the YIDD motif along with F358 and F360 of the thumb and V123 of RT2a (Figure 2E with wavy thin colored lines demarcating lightly shaded surface overlays). The contribution of I238 of the YIDD motif to the hydrophobic tunnel potentially results in a more stable interaction with the RT3a knot than an A residue at this position. These hydrophobic interactions also suggest why G2L4 RT has higher primer extension activity at lower salt concentrations^20^, which weaken hydrophobic interactions, making it easier for G2L4 RT to transition from an inactive to an active conformation upon binding a nucleic acid substrate.

### The RT3a knot stabilizes G2L4 RT and regulates its enzymatic activity

To investigate the function of the RT3a knot, we tested different combinations of 3 mutations (R190E, R191E, and I202E) in residues that were suggested by the apoenzyme structure to stabilize the RT3a knot and/or its interaction with the G2L4 RT active site (Figures 2B and 2C). For biochemical assays, wild-type and mutant G2L4 RTs were expressed with an N-terminal maltose-binding protein (MBP) tag that stabilizes the protein without interfering with biochemical activities^29^. All three mutant proteins were expressed at the same level as the wild-type protein (Figure S3A), and their gel filtration profiles showed a similar dimer peak (Figure 3A). Differential Scanning Fluorimetry (DSF) indicated that all three mutant proteins were stably folded but had lower melting temperatures than the wild-type protein (Figure 3B), indicating that RT3a interactions stabilize the G2L4 RT apoenzyme.

**Figure 3.**
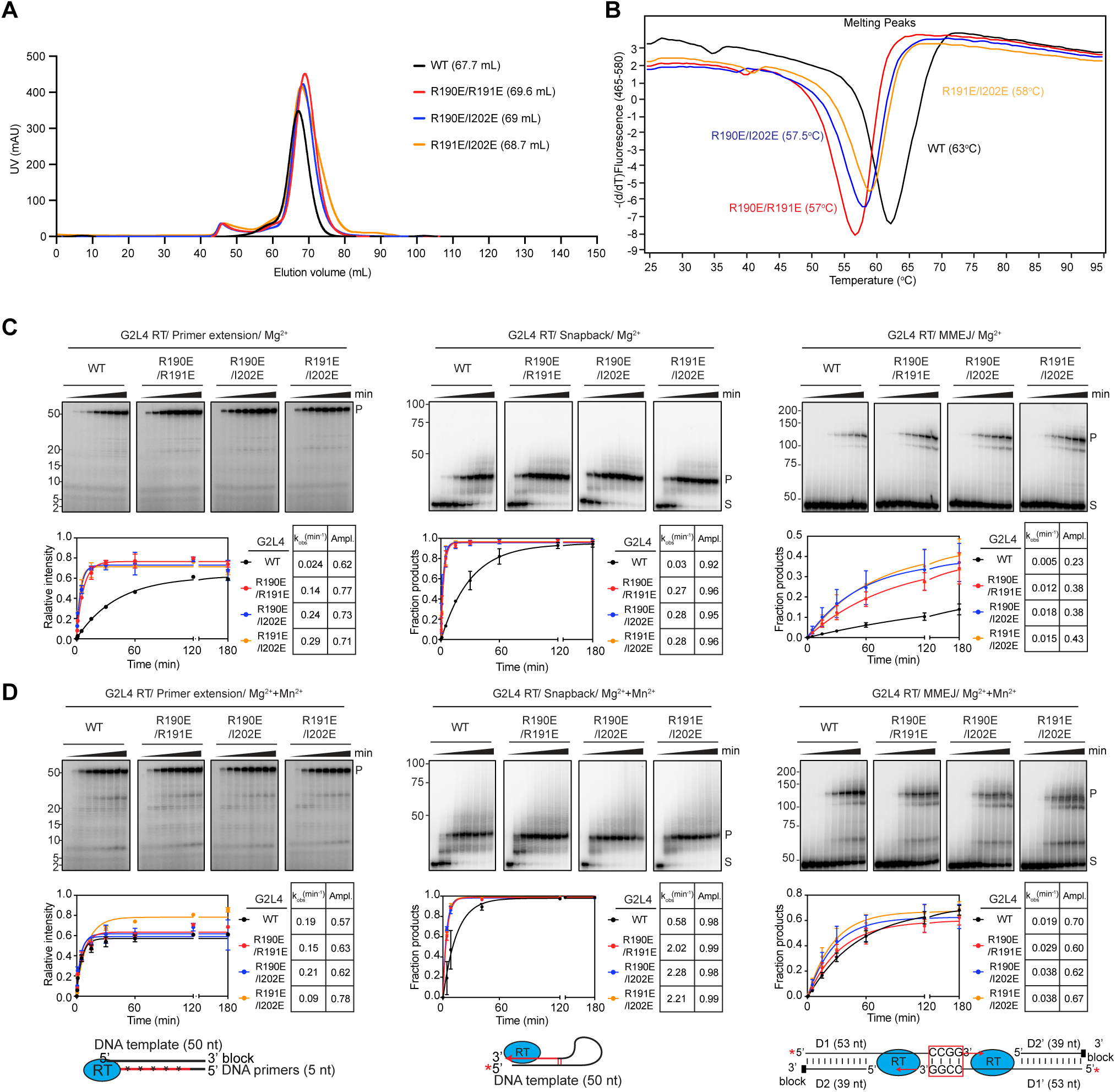
Biochemical analysis of the RT3a knot. (A) Size-exclusion chromatography of purified WT and RT3a mutant MBP-tagged G2L4 RTs. Elution profiles of different proteins are color-coded as indicated in the Figure. (B) Differential Scanning Fluorimetry (DSF) of WT and RT3a mutant MBP-tagged G2L4 RTs. Melting profiles of different protein are color-coded as in panel A. The plots show the derivative of fluorescence intensity as a function of temperature, indicating transitions or melting points indicative of structural changes or stability shifts. (C and D) Biochemical activities of WT and RT3a mutant MBP-G2L4 RTs in reaction media containing 10 mM Mg^2+^ (panel C) or 10 mM Mg^2+^ plus 1 mM Mn^2+^ (panel D). Left panels, primer extension assays with a 50-nt 3’-inverted dT-blocked DNA template, a 5-nt DNA primer, and ^32^P-labeled (red asterisk) dNTPs; middle panels, snapback DNA synthesis assays with a 5’- ^32^P-labeled 50-nt DNA substrate initiated by adding dNTPs; right panels, MMEJ assays with a 5’-[^32^P]-labeled 53-nt pre-annealed partially double-stranded DNAs with an inverted dT 3’- blocking groups on one strand and 3’ overhangs with complementary 3’ CCGG sequences on the opposite strand. The pre-annealed DNAs on the left side are denoted D1/D2, and those on the right side are denoted D1’/D2’. Primer extension reactions were initiated by adding an equimolar mix of 1 mM dNTPs (1 mM each of dATP, dCTP, dGTP and dTTP) and trace [α-^32^P]-dTTP. snapback DNA synthesis and MMEJ reactions with 5’-labeled substrates were initiated by adding an equimolar mix of 1 mM dNTPs. Reactions were incubated at 37°C for times up to 180 min, and Products (P) and substrates (S) were analyzed by electrophoresis in a denaturing polyacrylamide gel against 5’-[^32^P]-labeled synthetic ssDNA size markers for primer extension reactions and in a non-denaturing polyacrylamide gel against 5’-[^32^P]-labeled dsDNA size markers (Low Molecular Weight DNA Ladder; New England Biolabs) for snapback DNA synthesis and MMEJ assays. Size markers were run in parallel lanes with positions indicated to the left of the gels. The plots show the fraction of product based on the relative intensity of product and substrate bands as a function of time. The Tables to the right of the plots indicate the rate constant (k_obs_) and amplitude (Ampl.) for the production of products with curves fit to a first-order rate equation to obtain average values and variance indicated by error bars for two repeats of the experiment.

To explore how the RT3a knot affects the enzymatic activities of G2L4 RT, we compared the primer extension, snap-back DNA synthesis, MMEJ, and terminal transferase activities of wild-type (WT) G2L4 RT and the three RT3a mutant enzymes (Figures 3C, 3D, and S3B). The assays were done as time courses in reaction medium containing 10 mM MgCl_2_ in the absence or presence of 1 mM MnCl_2_, which enhances the DNA polymerase and strand annealing activities of WT G2L4 RT^20^. In the absence of Mn^2+^, all of the assayed biochemical activities were substantially higher for the three RT3a mutants than those for the wild-type enzyme, indicating that mutations that destabilize RT3a interactions with the active site make it easier for DNA substrates to displace the RT3a knot (Figure 3C). Addition of Mn^2+^ increased the activity of both the wild-type and mutant enzymes, but to a greater extent for the wild-type enzyme bringing its activity closer to that of the mutants, indicating that Mn^2+^ weakens the interaction of RT3a with the active site (Figures 3C, 3D and S3B). Collectively, these findings indicate that the RT3a knot is an evolutionary adaptation that stabilizes the G2L4 RT apoenzyme in an inactive configuration that impedes promiscuous interactions with cellular nucleic acids until it encounters a physiological substrate.

### Co-crystal structure of G2L4 RT bound to a snap-back DNA substrate

Although several cryo-EM structures of group II intron RTs bound to group II intron RNAs have been determined^4,30–36,40^, it has been difficult to obtain structures of group II intron RTs in an active conformation with bound template-primer and incoming dNTP, with GII RT being the only group II intron RT for which such a structure has been determined^4^. As an alternate approach for obtaining a structure of G2L4 RT in an active conformation, we returned to the previous finding that G2L4 RT has a strong strand-annealing activity that enables it to perform “snapback DNA synthesis”, a reaction in which the 3’ end of a DNA substrate folds back and initiates DNA synthesis by base pairing with a short stretch of complementary upstream nucleotides^20^.

Taking advantage of this activity, we obtained diffracting crystals of G2L4 RT bound to a 15-nt snapback DNA substrate (5’-AAGCGGTTAACCCAA-3’) that enabled initiation of DNA synthesis by base pairing of its two 3’ A residues (A14 and A15) to TT residues at upstream positions 7 and 8 (T7 and T8; Figure 4A). To stabilize these short base-pairing interactions, the co-crystallization set up included Mn^2+^, which favors strand annealing by G2L4 RT^20^, plus dCTP and dideoxy GTP (ddGTP) to enable addition of three complementary nucleotides at positions +1 to +3 ending with ddGTP and followed by base pairing of an incoming dCTP (position -1) opposite the templating base (G3 in the snapback substrate). The crystals had a rod morphology and diffracted to 2.77 Å.

**Figure 4.**
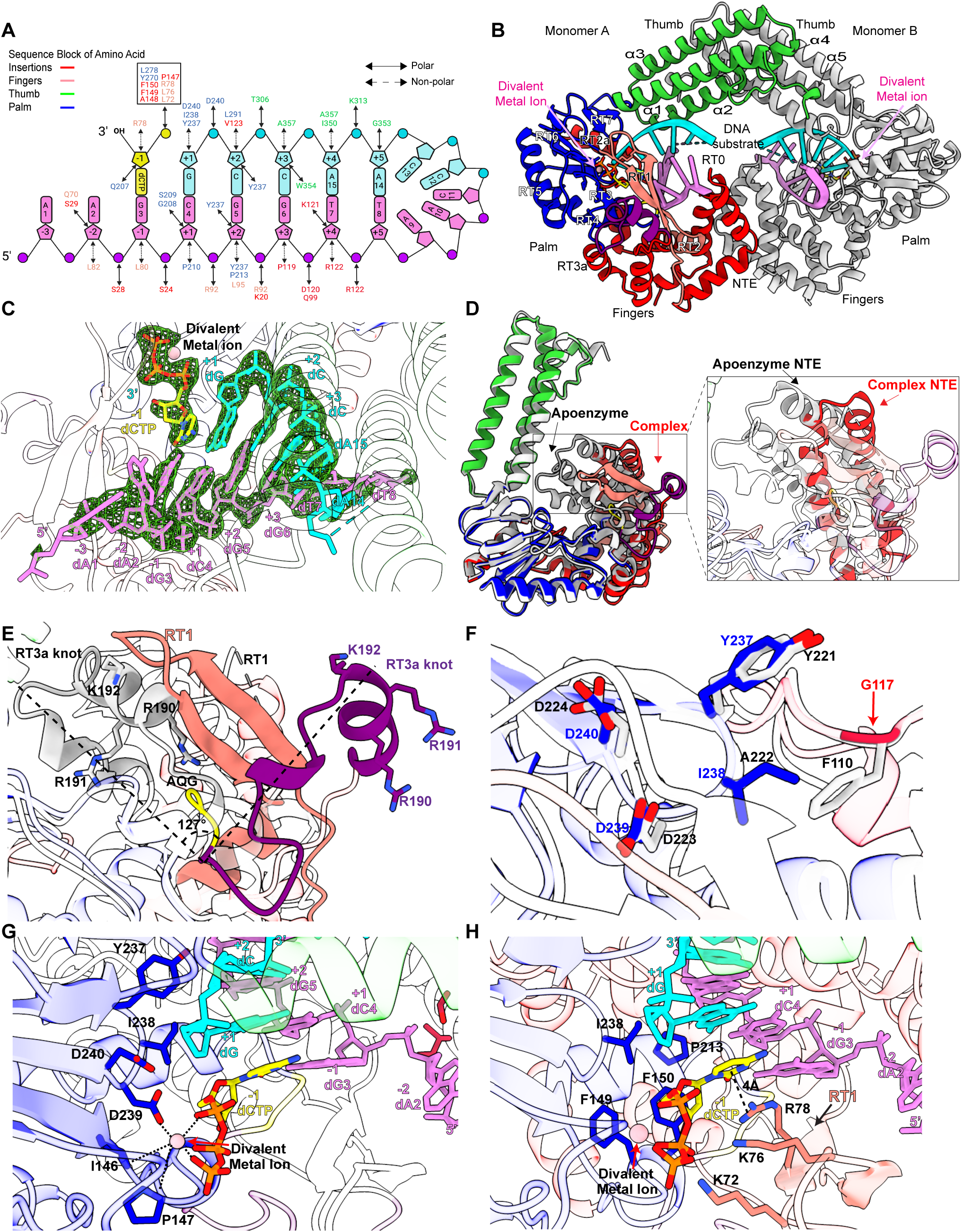
Structure of G2L4 RT in active conformation bound to a snapback DNA substrate and incoming dNTP. (A) Schematic of the snapback DNA synthesis substrate depicting its interactions with G2L4 RT amino acid residues identified in the crystal structure. Bases, ribose rings, and phosphates are represented by rectangles, pentagons, and circles, respectively. Interactions between amino acids and nucleotides are indicated by double-arrowheaded black lines for polar interactions and double-arrowheaded dashed lines for non-polar interaction. Amino acid residues that interact with the snapback substrate or incoming dCTP are color-coded by their location within the RT as indicated at the top left. Numbers within rectangles represent the sequence order of nucleotides in the substrate, with N-3 to N+5 corresponding to nucleotide positions relative to 5’ end of the template, and N-1corresponding to the templating nucleotide and incoming dCTP. Template strand nucleotides are colored violet; primer strand nucleotides, including those incorporated from the crystallization mix, are colored cyan; and the incoming dCTP at the active site is colored yellow. (B) Structure of G2L4 RT snapback DNA synthesis complex. Monomer A regions are colored coded as in Figure 1A, and monomer B is colored gray. Divalent metal ions are shown as pink spheres with pink arrows pointing to their locations. RT3a is now on the outer surface of the protein. (C) A Fo-Fc map calculated by subtracting the calculated electron density (Fc) from the observed electron density (Fo) for G2L4 RT to identify differences due to binding of the snapback DNA substrates at the active sites. Positive density of the electron cloud is shown as green mesh contoured at a level of 3 sigma (σ), with different regions of the snapback DNA substrate colored as in panel A. (D) Superimposition of a G2L4 RT monomer in snapback DNA complex with regions colored-coded as in Figure 1A and a monomer of the G2L4 RT apoenzyme colored gray. The two monomers are aligned at the palm. The right panel shows a close-up of the NTE region in the snapback DNA complex structure compared to that in the apoenzyme structure. (E) Superimposition of G2L4 RT snapback complex and apoenzyme structures showing differences in RT1 and the RT3a knot. RT1 is colored salmon, the AQG sequence at the beginning of RT4 yellow, and the RT3a knot purple. The dashed black lines depict the shift in position of the RT3a knot in the G2L4 RT apoenzyme structure compared to that in the snapback complex structure. Charged residues RRK (190-192) in the RT3a knot of the snapback complex structure are shown as purple sticks. (F) Superimposition of the active site of G2L4 RT in the snapback complex (lightly colored-coded as in Figure 1A) and GII RT (gray). Side chains of key active-site residues in G2L4 RT are shown as sticks labeled and color-coded by region as in Figure 1A, and those in GII RT are shown in gray and labeled in black. Bound nucleic acids are not shown. (G) Active-site interactions of G2L4 RT with incoming dCTP and DNA substrate. A divalent metal ion (pink sphere) di-pyramidally coordinated with active site residues (blue, sticks) and the incoming dCTP (yellow, stick) are depicted by dashed black lines. Nucleotides in the snapback complex structure are numbered and color-coded as in Figure 4A. (H) Interactions between the incoming dCTP (yellow, stick) and RT1 motif. Charged residues K72, K76, R78 and hydrophobic residue F150 are shown as sticks and color-coded as Figure 1A. The connection between R78 and the ribose ring of the incoming dNTP is indicated by a black dashed line.

Initial attempts to solve the snapback structure by molecular replacement using the G2L4 RT apoenzyme structure as a search model failed, likely because of the different relative orientations of the fingers and palm/thumb in the apoenzyme and substrate-bound G2L4 RT structures. Instead, we defined the fingers and palm/thumb as separate rigid bodies during molecular replacement to allow for mobility between these two protein regions. This led to a solution in space group P2_1_2_1_2_1_ with the two G2L4 RT monomers bound to two separate snapback DNA substrates per asymmetric unit (Figure 4B). A Fo-Fc map that showed positive density (green) in areas where an initial model lacked atoms guided us to fit the snapback DNAs into the correct position (Figure 4C). After fitting the snapback DNA substrates into the model based on the Fo-Fc map, we used a 2Fo-Fc map to further validate the model (Figure S4A). This confirmed that the newly placed DNA atoms fit well into the overall electron density in a way consistent with the experimental data. The final model corresponds to the full-length G2L4 RT protein with the DNA ligands fitted into the electron density that accounts for the snapback DNAs (Figures 4C and S4A). The two monomers were again virtually identical with RMSD of 0.78 Å.

As expected, the structure showed that A14 and A15 at the 3’ end of the snapback DNA substrate annealed to the upstream residues T8 and T7, respectively (Figures 4A and 4C). The intervening AACCC loop (DNA substrate nucleotides 9-13) was not visible in the structure and presumably disordered. Because dCTP and ddGTP were included in the crystallization solution, two C residues complementary to G5 and G6 and ddGTP complementary to C4 of the template were added to the 3’ end of the DNA substrate, stabilizing the annealed microhomology (Figure 4C). As hoped, dCTP complementary to G3 was bound at the active site but not incorporated due to the lack of free hydroxyl of the preceding 3’-terminal dideoxyguanosine (Figures 4A and 4C).

Compared to its clutched apoenzyme structure, each G2L4 RT monomer adopted an active configuration with the NTE moving outward to accommodate the bound DNA (Figure 4D) and with the bound DNA substrate pushing the RT3a knot completely out of the active site with a rotation of 127° from its position in the G2L4 apoenzyme structure (Figure 4E). Extrusion of the RT3a knot allowed the disordered RT1 in the apoenzyme structure to form anti-parallel β-strands over the active-site pocket as in other RTs and the AQG motif to revert to its functional position to coordinate alignment of the incoming dNTP, resulting in an active structure similar to that of GII RT (Figures 4D, S4B, and S4C)^4^. Except for a short two-turn α-helix (residue 186-193), the extruded RT3a knot became almost completely random coil, indicating that its interaction with the active site stabilized its more complex secondary and tertiary structures in the apoenzyme (Figure 4E). Notably, three conserved basic residues of the RT3a knot (R190, R191, and K192), which interacted with the active site in the apoenzyme structure, were now exposed to bulk solvent, where they might serve as initial contact points for sampling potential nucleic acid substrates (Figure 4E).

### Configuration of the active site with bound DNA substrate and incoming dNTP

Although the overall active structures of G2L4 and GII RTs are similar, the snapback DNA synthesis structure revealed a series of structural adaptations that optimize the DSBR activity of G2L4 RT. These include the conserved YIDD motif instead of YADD at the active site, a distinguishing conserved feature of G2L4 RTs, with I favoring strand annealing in G2L4 RT and disfavoring primer extension in both G2L4 and GII RT^20^. Except for the I/A substitution with compensation and consequences discussed further below, the configuration of the YIDD motif and its interactions with substrates closely resembled those of the YADD motif in GII RT (Figures 4A, 4F, S4B, and S4D)^4^. The Y237 side chain hydroxyl group hydrogen bonds with the nucleotide base of G5, and the first D (D239) of the YIDD motif coordinates a divalent metal ion di-pyramidically along with the phosphate groups of incoming dCTP (Figure 4G). D240 faces outwards away from the active site as seen in previous structures of GII RT and HIV RT^4,25,41^.

The reformed RT1 lid over the active site stabilizes the triphosphate group of the incoming nucleotide by salt bridges with positively charged residues K72, K76, and R78 (Figure 4H). Notably, the side chain of R78 aligns parallel with the terminal base of the substrate (ddG+1) via a cation-π interaction of 4 Å, close to the base-pair distance of 3.4 Å. This favorable interaction likely stabilizes the binding of the incoming base and may act as a bookend to push the dNTP forward into the active site aligned with the preceding base pairs (Figure 4H). The R78 residue in G2L4 RT corresponds to a conserved arginine residue that adopts the same configuration and likely plays a similar role in GII RT (R75), HIV-1 RT (R72), and HCV RdRP (R158) (Figures S1C and S4D)^4,25,42^. Further stabilization of the incoming dNTP in G2L4 RT comes from F150 of RT3, which interacts hydrophobically with the ribose sugar pucker of the incoming dCTP (Figure 4H). This phenylalanine, which is also conserved in group II intron RTs (F143 in GII RT; Figure S1C)^4^ and replaced by a tyrosine in HIV-1 RT^26^, may play a role in stabilizing the incoming nucleotide during polymerization.

The conserved active site I residue (I238) in G2L4 RT, which favors strand annealing^20^, potentially stabilizes the active configuration of the enzyme by fitting into a hydrophobic pocket formed by F149/150 and P213 with its side chain stacking hydrophobically with the sugar pucker of the base of the incoming dCTP, a stronger hydrophobic interaction than that for the conserved A residue in the YADD motif of GII RT (Figure 4H). The previous finding that I for A substitution in GII RT decreased the rate of primer extension^20^ is likely due to Van der Waals clashes of the substituted I residue with F110, which extends into the GII RT active site from RT2a (Figures 4F and S1C)^4^. In G2L4 RT, however, this phenylalanine residue in RT2a is replaced by a conserved Gly residue (G117), a covariation that leaves enough space for the Ile side chain to fit into the active site without steric hinderance (Figures 4F and S1C). The substitutions at the active site that enhance the DNA repair function of G2L4 RT may come at the expense of lower fidelity, a trade-off of human DNA polymerase θ and other DNA repair polymerases^43,44^.

### RT0-loop and thumb domain interactions that favor annealing of microhomologies

The RT0 loop at the end of the NTE is a key structural feature of non-LTR-retroelement RTs that is required for the MMEJ activity but not the primer extension activity of both G2L4 and GII RTs^20^. In the crystal structure of GII RT with a bound RNA template-DNA primer, the NTE inserts into the major groove of the template-primer duplex with the RT0 loop forming an 8-aa lid over the active site between positions -1 to +1 (Figures 5A and 5B)^4^. Additionally, two tyrosine residues in thumb helix a2 (Y318 and Y325) stabilize alignment of the DNA primer with the RNA template via phosphate backbone hydrogen bond and π-π stacking interactions, respectively (Figures 5A and 5C)^4^.

**Figure 5.**
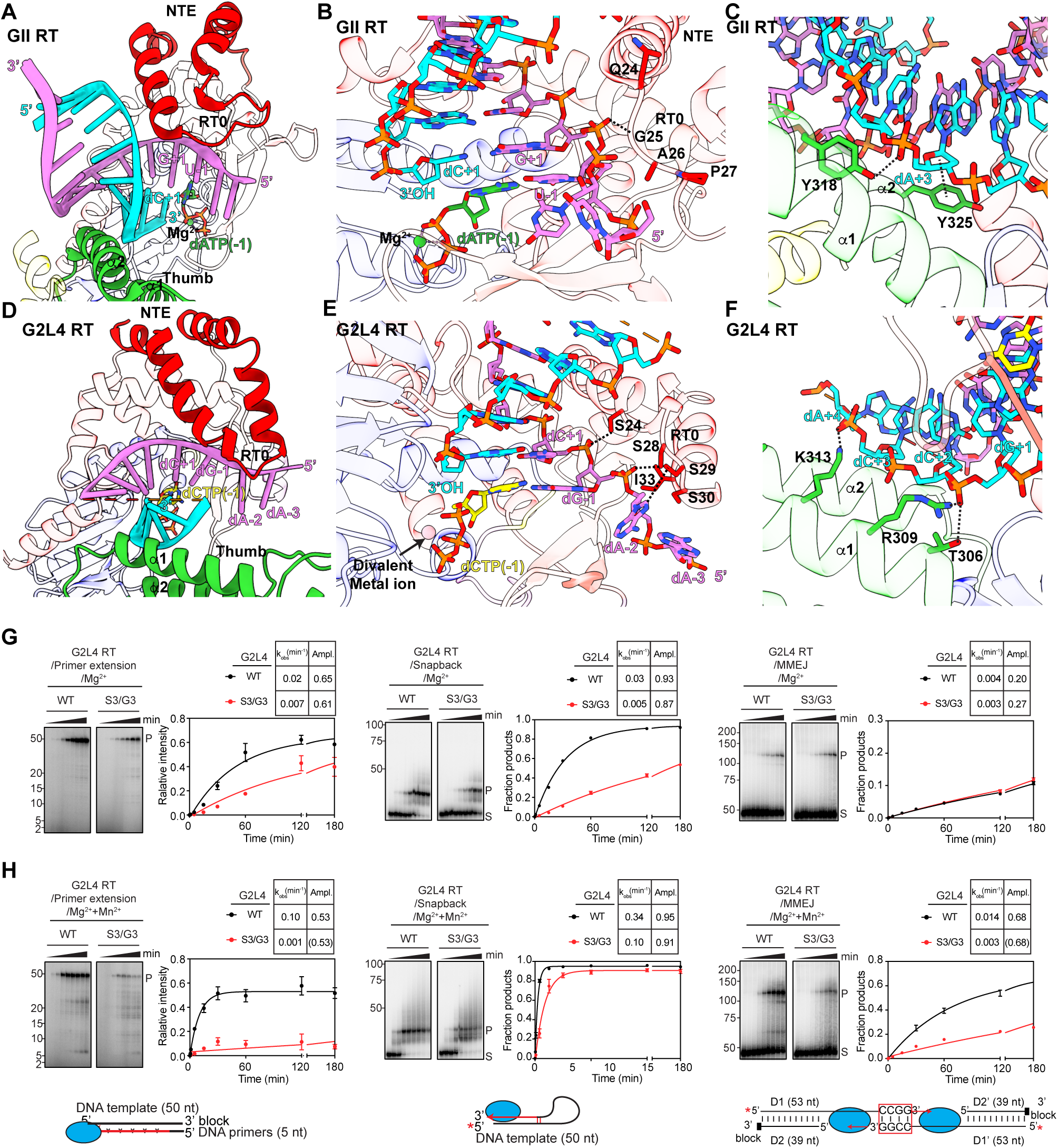
RT0-loop and thumb domain interactions with the annealed microhomology in the snapback DNA complex structure. (A) Wide-angle view of NTE/RT0 (red) and thumb domain (green) interactions in a GII RT complex structure^4^ with RNA template (violet), annealed DNA primer (cyan), and incoming dATP (green) shown as sticks, and a catalytic Mg^2+^ shown as a green sphere. Numbers indicate nucleotide positions relative to 5’ end of the template, with -1 corresponding to the position of the templating nucleotide and incoming dATP. (B) Close-up view of interacting (dashed black line) residues of GII RT RT0 loop and the RNA template. (C) Close-up view of interacting residues of GII RT thumb domain and DNA primer. (D) Wide-angle view of the NTE/RT0 (red) and thumb domain (green) interactions in the G2L4 RT snapback DNA complex structure with the DNA template strand (violet), annealed to DNA primer strand (cyan), and incoming dCTP (yellow) depicted as sticks. Numbers indicate nucleotide positions relative to the 5’ end of the template, with N-1 corresponding to the position of the templating nucleotide and incoming dCTP. (E) Close-up view of the interacting serine residues of G2L4 RT NTE/RT0 loop and DNA template strand in the snapback DNA complex structure. A divalent metal ion is shown as a pink sphere. (F) Close-up view of the interacting residues of G2L4 RT thumb domain and DNA primer strand in the snapback DNA complex. (G and H) Biochemical assays of WT and S3/G3 mutant G2L4 RTs in reaction media containing 10 mM Mg^2+^ (panel G) or 10 mM Mg^2+^ plus 1 mM Mn^2+^ (panel H). Left panel, primer extension assays; middle panels, snapback DNA synthesis assays; right panel, MMEJ assays. Biochemical assays were done as in Figures 3C and 3D. The plots show the average values and variance for two repeats of the experiment. Red asterisks in schematics at the bottom indicate an internally labeled DNA strand for primer extension assays or a 5’-labeled strand for snapback DNA synthesis and MMEJ assays.

For comparisons with GII RT, we focused on the corresponding regions of the G2L4 RT snapback DNA structure in which the 3’ end of the DNA served as a primer strand that initiated DNA synthesis by annealing to a short (2 nt) microhomology within an upstream region of the DNA substrate that served as the template strand. These comparisons showed that the longer NTE of G2L4 RT extended farther out over the template strand with its smaller RT0 loop (5 aa in both the snap back and apoenzyme structures) binding directly to the phosphate backbone between positions -2 and +1 (Figures 5D, 5E, and S1C). The alignment of the DNA primer strand at the annealed microhomology differed from GII RT in being stabilized via hydrogen bond and salt bridge interactions with T306, R309 and K313 at the bottom of the thumb helix a1 (Figures 5D and 5F).

A further adaptation of the NTE/RT0 loop for the DSBR function of G2L4 RT was a set of four serine residues (S24 in the NTE and S28-S30 at the center of the RT0 loop) that were strongly conserved in G2L4 RTs, but not found in GII or other group II intron or non-LTR-retro-element RTs (Figures S1C and S5A)^20^. The snapback DNA synthesis structure indicated that hydroxyl side chains of these serine residues play a major role in direct binding to the template strand (Figure 5D and 5E). S28 hydrogen bonds to the backbone phosphate between dG-1 and dA-2, while S29 hydrogen bonds to the dA-2 base (Figure 5E). S30, while not directly interacting with the template strand, forms an intramolecular hydrogen bond with the backbone nitrogen of I33 to stabilize a β-turn that protrudes S28 and S29 to interact with the template strand (Figure 5E). S24 in the NTE forms an H-bond with the phosphate backbone between dG-1 and dC+1, further stabilizing the extended interaction between the NTE/RT0 loop and the template strand (Figure 5E).

To assess the contribution of the three consecutive conserved serine residues (S28-S30) in the RT0 loop, we constructed a G2L4 RT mutant in which all three of these residues were changed to Gly (S3/G3). Biochemical assays showed that the substitution of Gly at these positions in the RT0 loop inhibited most biochemical activities of G2L4 RT: primer extension (∼3-fold in the absence of Mn^2+^ and 100-fold in the presence of Mn^2+^); snapback DNA synthesis (6-fold in the absence of Mn^2+^ and 3-fold in the presence of Mn^2+^); MMEJ (roughly equal in the absence of Mn^2+^ and 4-fold lower in the presence of Mn^2+^; Figures 5G and 5H) with similar results for terminal transferase assays with different dNTPs (Figure S5B). These findings suggest that the conserved RT0 loop structure and substrate contacts of the conserved Ser residues contribute to diverse biochemical activities. Collectively, these findings indicate that the NTE/RT0 loop of G2L4 RTs evolved to play a critical role in MMEJ that differs mechanistically from that of GII and other non-LTR-retroelement RTs (see Discussion).

### Mutations that weaken dimerization affect enzymatic activities of G2L4 RT

The G2L4 dimer interface in both the apoenzyme and co-crystal structures involve similar interactions (ionic, H-bond, and hydrophobic) between the outer surfaces of the NTE and thumb of the two monomers, with the thumb domains contributing the larger area to the dimer interface (Figures 6A-6C). This large thumb domain interface reflects in part that the α-helical bundle comprising the thumb of G2L4 RT extends straight up 15Å farther than in group IIC intron RTs in which a shorter α-helical bundle interacts with the DNA-binding domain to form a basic cleft that binds to a DNA hairpin structure at intron insertion sites (Figures 1C and 1D)^4,34,45^. The G2L4 RT thumb also has a 22-aa C-terminal extension (residues 390-411) that includes two short α-helices (α4 residues 394-401 and α5 residues 407-411) that interact with the thumb domain of the opposite monomer (Figure 1B, 1C, and S1C). We confirmed that G2L4 RT remains a dimer in the absence of the N-terminal MBP tag, although removal of the tag decreased the thermostability of the protein measured by DSF, reflecting a stabilizing effect of the MBP tag on the overall structure of the protein (Figures S6A and S6B). Deletion of C-terminal residues 401 to 411, which include α5, likewise decreased protein stability, but did not prevent dimer formation (Figures S6A and S6B).

**Figure 6.**
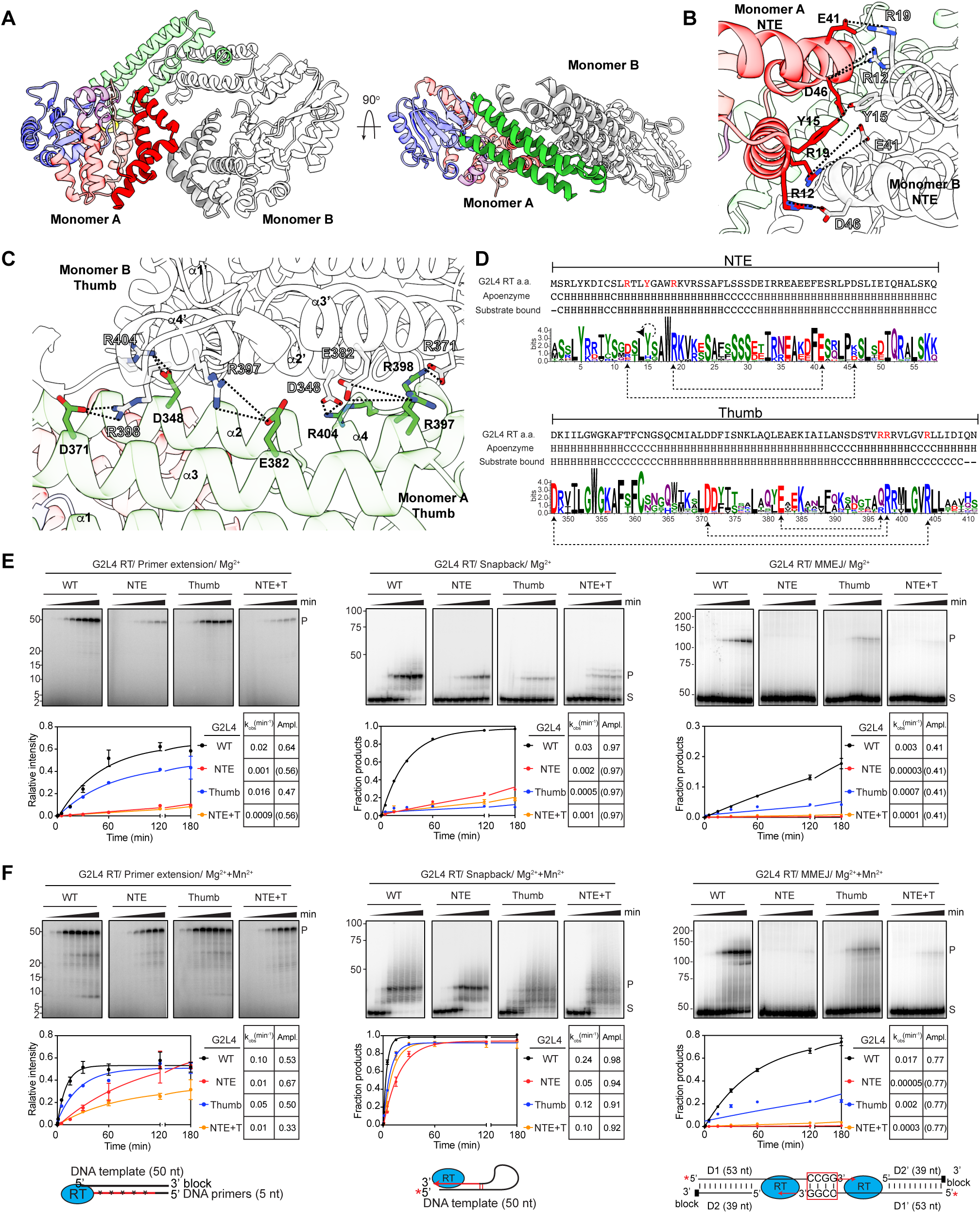
Structural and mutational analysis of interactions at the G2L4 RT dimer interface. (A) Left panel, view of a G2L4 RT dimer showing interacting regions of the NTEs of monomers A and B; right panel, 90° vertically rotated view showing interacting regions of the thumb domains of monomers A and B. Regions in the left-hand monomer are color-coded as in Figure 1A. (B) Interactions of NTE residues R12, Y15 and R19 of monomer A with NTE residues D46, Y15 and E41 of monomer B. The NTEs of monomer A and B are shown in red and white, respectively. Black dashed lines highlight salt bridges or hydrogen bonds. (C) Interactions of thumb domain residues R397, R398, and R404 of monomer A with thumb domain residues E382, D371 and D348 of monomer B. The Thumb domains of monomer A and B are shown in light green and white, respectively. Black dashed lines highlight salt bridges or hydrogen bonds. (D) WebLogos of interacting regions of the NTEs and the thumb domains of G2L4 RT dimers. Interactions between amino acid residues in the two monomers are indicated by black dashed lines with dashed circular line above the Y15 indicating interactions between the same residue in the two monomers. Amino acids in α-helical (H) and random coil (C) regions of apoenzyme and snapback substrate bound G2L4 RTs based on the crystal structures are indicated below the amino acid sequence. (E and F) Biochemical assays of G2L4 RT dimer interface mutants in reaction media containing 10 mM Mg^2+^ (panel E) or 10 mM Mg^2+^ plus 1 mM Mn^2+^ (panel F). Left panel, primer extension assay; middle panels, snapback DNA synthesis assays; right panel, MMEJ assays. Reactions were done as in Figures 3C and 3D. Ampl. values in parentheses indicate fixed amplitudes for reactions that did not reach an end point based on the average amplitude value for those that reached a clear end point during the experiment. The plots show the average values and variance for two repeats of the experiment. Red asterisks in schematics at the bottom indicate an internally labeled DNA strand for primer extension assays or a 5’-labeled strand for snapback DNA synthesis and MMEJ assays.

To test if dimerization of G2L4 RT is functionally important, we constructed three G2L4 RT mutants in which NTE and/or thumb residues identified as contributing to interactions that potentially stabilize the dimer interface were replaced by alanines: (i) NTE residues R12, Y15 and R19 (denoted NTE); (ii) Thumb residues R397 and R398 in α4 and R404 in a random coil region between α4 and α5 (denoted Thumb); and (iii) NTE + Thumb residues R12, Y15 and R398 (denoted NTE + T; Figures 6B-6D). Analysis of the mutant proteins on a non-denaturing gel showed that the Thumb and NTE + T mutant proteins had higher proportions of G2L4 RT monomers (25% and 36%, respectively) than wild-type G2L4 RT, while the NTE mutant protein ran entirely as dimers, as did all other G2L4 RT mutants analyzed in this study (Figure S6C). All three of the dimer interface mutants had decreased thermostability, with the NTE + T and Thumb mutants having lower melting temperature than the NTE mutant (Figure S6B). These findings indicate that dimerization contributes to the stability of G2L4 apoenzymes and that the tested thumb domain interactions involving α4 and the random coil region make a stronger contribution to dimer formation and protein stability than the tested NTE interactions.

Biochemical assays showed that three dimer interface mutants generally had lower primer extension, snap-back DNA synthesis, MMEJ, and terminal transferase activities than did the WT enzyme in the absence or presence of Mn^2+^, but surprisingly with these decreases more pronounced for the NTE and NTE+T mutants than for the Thumb mutant, which had nearly equal or slightly higher activity than the WT enzyme in some assays (Figures 6E, 6F and S6D). While all three dimer interface mutants had substantial primer-extension activity and close to WT snapback DNA synthesis activity in the presence of Mn^2+^, the NTE and NTE+T mutants had little if any MMEJ activity in the presence or absence of Mn^2+^ (Figure 6E and 6F). Collectively, these findings indicate that mutations of interacting residues at the dimer interface affect the overall stability and biochemical activities of G2L4 RT, but with the NTE mutations, which have no detectable effect on dimerization, having a disproportionally large effect on MMEJ compared to other biochemical activities.

### Model for MMEJ using both monomer subunits of G2L4 RT

The snapback DNA synthesis structure revealed a mechanism potentially analogous to that used for MMEJ in which a G2L4 RT dimer bound to the 3’ end of a single-stranded DNA ending with 3’ AA residues (primer strand) and searched upstream regions of the single-stranded DNA (template strand) for complementary TT residues to which the 3’ end could base pair for initiation of DNA synthesis (Figure 4). At the high concentrations of the short snapback substrate used to obtain crystals, both monomers had activated structures bound to separate short snapback DNA substrates, but we were unable to obtain crystals with longer snapback DNA substrates that could bind across both monomers, likely due to steric hindrance for reasons described below. Despite conformational changes during substrate binding and initiation of DNA synthesis coupled to reconfiguration of RT3a, the dimer interface remained identical for both the apoenzyme and activated forms of G2L4 RT (Figure 7A). This finding indicated that G2L4 RT could remain a stable dimer with one monomer activated and the other remaining inactive.

**Figure 7.**
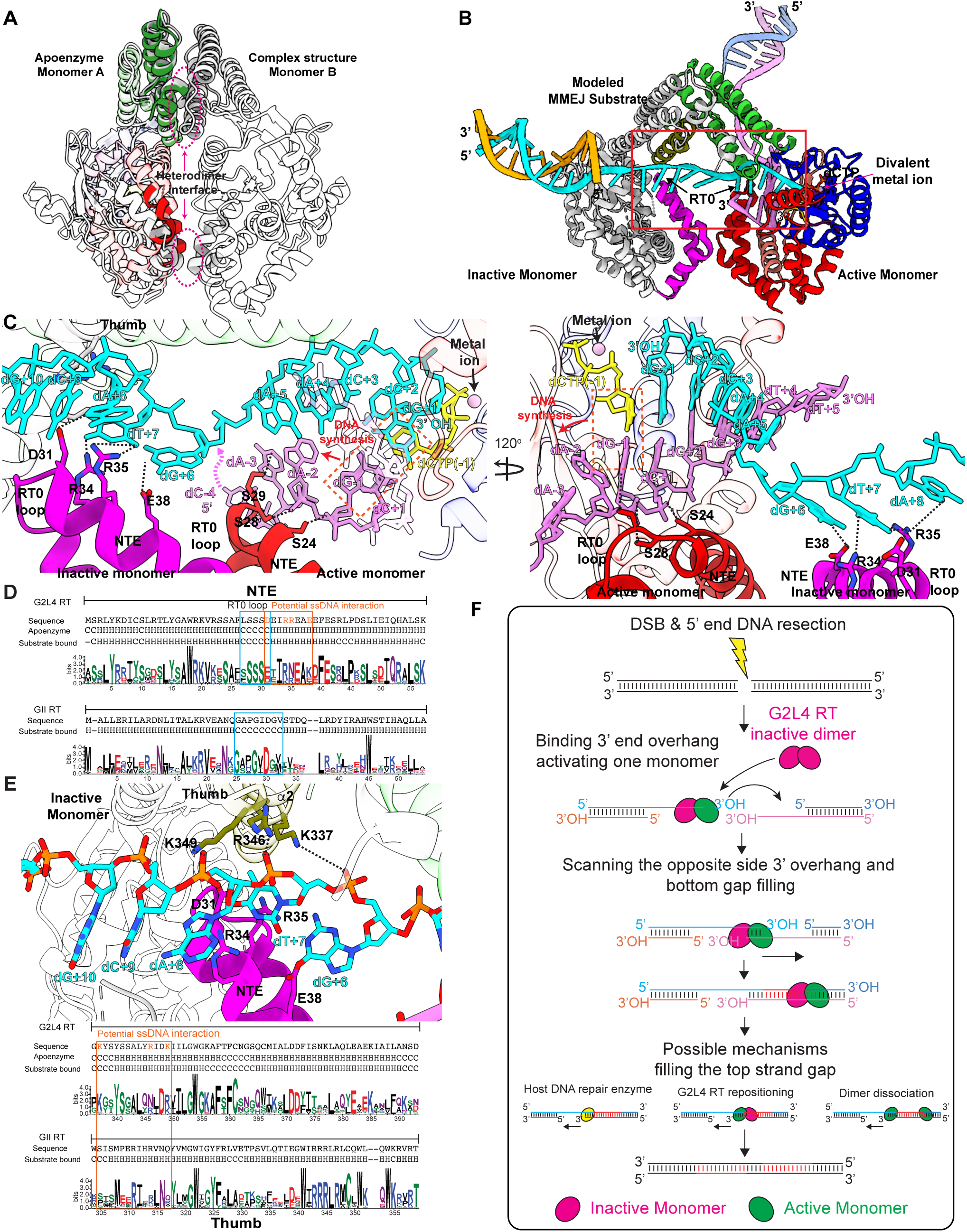
Model of the G2L4 RT MMEJ mechanism. (A) Superimposition of the G2L4 RT apoenzyme structure (NTE and thumb regions colored red and green, respectively) and snapback DNA synthesis structure (gray/white) aligned at monomer B. The dimer interfaces between the NTE and thumb regions are encircled by magenta dashed lines. (B) Model of a G2L4 RT dimer comprised of an activated monomer (right) and a trailing inactive monomer (left) bound to the 3’ end of a single-stranded 3’-DNA overhang from the left side of a DSB (cyan) annealed to a 5-bp microhomology at the 3’ end of the 3’-overhang from the right side of the DSB (violet). Regions of the active monomer (right) are colored as Figure 1A, and the NTE/RT0 loop and the a2 helix of the thumb domain of the inactive monomer (left) are colored purple and olive, respectively, with other regions gray. Regions of the protein interacting with the MMEJ substrate are highlighted in a red box. (C) Close-up 120° horizontally rotated views of panel B highlighting interactions of the NTE/RT0 loops of the leading active monomer and trailing inactive monomer relative to the annealed microhomology between single-strand 3’-overhangs from the left (cyan) and right (violet) sides of the DSB. The interactions (dashed black lines) of serine residues in the NTE/RT0 loop of the active monomer with dA-2 to dC+1 of the 3’-overhang from the right side of the DSB (violet) are based on the snapback DNA synthesis structure in Figure 5D and 5E, and potential interactions of the inactive monomer NTE/RT0 loop and thumb domain with the 3’- overhang from the left side of the DSB (cyan) follow from positioning of the active monomer with no further adjustments. Base pairing of the incoming dCTP with dG-1 of the template strand (violet) is highlight in an orange dashed line box. The direction of DNA synthesis filling in the single-stranded 3’overhang on the right side of the DSB is indicated by a red arrow. Regions upstream of dC-4 of the 3’-overhang from the right side of the DSB (dashed violet line) are not shown. (D) WebLogos comparing the NTE/RT0 loop regions of G2L4 RTs to those of GII RTs. Amino acid residues of the trailing inactive G2L4 RT monomer that potentially interact with the single-stranded DNA dG+6 and dA+8 in the gap between the two monomers on the left side of the DSB are highlighted in orange. α-helical (H) and random coil (C) regions in the inactive apoenzyme and active snapback DNA synthesis structures of G2L4 RT and the template-primer complex structure of GII RT^4^ are indicated below the amino acid sequences. (E) Model of interactions between amino acid residues in the thumb domain of the inactive monomer and the single-stranded gap between inactive and active monomers in 3’-overhang from the left side of a DSB. WebLogos below the model compare thumb domain regions of G2L4 RTs that potentially interact with the single-stranded gap to those of GII RTs, with potentially interacting residues in G2L4 RT highlighted in orange. α-helical and random coil regions based on crystal structures of G2L4 and GII RTs are indicated below the sequences. (F) Schematic of the suggested mechanism by which G2L4 RT functions in DSBR by MMEJ.

Under physiological conditions where DNA substrate binding is likely rate-limiting, activation of both monomers by binding to the 3’ end of a single-strand DNA overhang at a DSB site likely occurs sequentially if at all. Utilizing the snapback complex and apoenzyme structures, we modeled a G2L4 RT dimer bound to the 3’ end of the 3’-overhang from the left side of a DSB (primer strand, cyan) annealed to a 5-bp microhomology at the 3’ end of the 3’-overhang from the right side of the DSB (template strand, violet; Figure 7A-E; schematic Figure 7F). In this model, the leading monomer (right, regions color-coded as in Figure 1A) is activated by binding the 3’ end of the primer strand (cyan), while the trailing monomer (left, gray with NTE/RT0 loop magenta and thumb α2-helix olive) remains inactive (Figure 7B). The model, based entirely on the two structures with no further adjustments, showed that the NTE/RT0 loops of both monomers were positioned by their interactions at the dimer interface on opposite sides of the annealed 5-bp microhomology, with the leading active monomer positioned to copy the 3’-overhang on the right side of the DSB (Figure 7B and 7F),

In close-up views of the model (Figure 7C), the NTE/RT0 loop of the leading active monomer bound to the single-strand 3’-overhang on the left side of the DSB (primer strand, cyan) is positioned over the annealed microhomology to bind the single-strand 3’-overhang on the right side of the DSB (template strand, violet) via phosphate backbone interactions at dA-2 to dC+1 using conserved serine residues as in the snapback DNA complex structure (see Figure 5D and E). With this position fixed, NTE/RT0 of the trailing inactive monomer is positioned to interact with three exposed bases (dG+6 to dA+8) of the priming strand (cyan) in the gap between the two monomers via hydrogen bonds from residues D31, R34, R35, and E38 (Figure 7C). Supporting these interactions: (i) charged residues at these positions in the NTE were conserved in G2L4 RTs, while only the cognate of D31 (D30 in GII RT) was conserved in the larger RT0 loop of GII RTs (Figures 7C, 7D and S1C), and (ii) mutations in which R35 by itself or together with R34 or E38 were replaced by alanines preferentially inhibited MMEJ (Figures S7A and S7B). Combined R34A/D31A mutations had little or no effect on MMEJ or primer extension *in vitr*o, suggesting a smaller or indirect contribution of these residues (Figures S7A and S7B). A critical role for the NTE interactions at the dimer interface that position the NTE/RT0 loops of both monomers on an MMEJ substrate is supported by the finding that mutations that affect these interactions had no detectable effect on dimerization (Figure S6A), but almost totally abolished MMEJ, even in the presence of Mn^2+^ which stimulated other biochemical activities (Figures 6E and 6F).

At an earlier stage of MMEJ, the binding of a G2L4 RT dimer to the single-stranded 3’- overhang on the left side of the DSB (priming strand) may stabilize it in an extended conformation that facilitates scanning the 3’-overhang on the opposite side of the DSB for annealing to a suitable microhomology. In addition to the above interactions, the model with no further adjustments suggests that the α2-helix of the thumb domain of the inactive monomer (olive) may contribute to such a function by phosphate-backbone interactions of K337, R346, and K349 with dG+6, dT+7 and dA+8 of the 3’-overhang from the left side of the DSB (Figure 7E). These amino acid residues, which do not contribute to the dimer interface, were conserved as positively charged residues in G2L4 RTs but not GII RTs (Figures 7E and S1C). A similar model in which both monomer subunits were in an active conformation displaced the RT0 loop of the trailing monomer away for the MMEJ substrate (Figure S7C), and a model with an active leading and trailing inactive monomer and annealed microhomology >6 bp showed a steric clash between the 7th base pair and the α5-helix of the thumb domain of the inactive monomer (Figure S7D), consistent with previous findings that G2L4 RT prefers MMEJ substrates with shorter annealed microhomologies and longer single-strand gaps than GII RT, which functions as a monomer^20^.

The model for the initial steps of MMEJ by G2L4 RT positions the enzyme to fill in the single-stranded gap on only the right side of the DSB (Figure 7B and 7F), potentially critical for cell viability by generating a continuous single-strand DNA across what was an annealed microhomology between the two 3’-overhangs at a DSB. The remaining single-stranded gap on the left side of the DSB could be filled in by several possible mechanisms, including (i) activation of the trailing monomer after DNA synthesis to the right by binding of 3’ end of the single-strand 3’ DNA overhang from the right side of the DSB followed by monomer dissociation and continued DNA synthesis by each monomer in opposite directions; (ii) repositioning of the first partially activated dimer or binding of second partially activated dimer; or (iii) by cellular DNA repair enzymes, which are also needed to fully repair the DSB (Figure 7F).

## Discussion

Here, we obtained crystal structures of a chromosomally encoded G2L4 RT apoenzyme and its snapback DNA synthesis complex, revealing features that evolved to optimize its cellular function in DSBR via MMEJ and suggesting a model for the MMEJ mechanism shown in Figure 7F. After synthesis, the G2L4 RT apoenzyme forms a homodimer with each subunit stabilized by the RT3a knot, a novel structural feature of G2L4 RTs that was not predicted by the most recent version of AlphaFold 3.0^39^. The G2L4 RT apoenzyme structure showed that the RT3a knot completely blocks the active site of each monomer, replacing catalytic divalent metal ions and rendering the enzyme totally inactive until it binds a physiological substrate (Figures 2C and 2D). Mutations that weakened the interaction of the RT3a knot with the RT active site led to higher biochemical activities across all tested substrates, suggesting an equilibrium between an inactive and an activatable conformation that can lead to full binding of physiological substrates (Figures 3 and S3). In the activatable conformation, conserved basic residues of RT3a that block the active site of G2L4 RT move to the surface of the protein, potentially enabling sampling of nucleic acid substrates by non-specific binding.

The snapback DNA synthesis structure indicated a mechanism likely relevant to MMEJ in which a G2L4 RT dimer binds the 3’ end of a single-stranded DNA substrate that serves as a primer and searches for complementary microhomologies in upstream regions of the snapback substrate that serves as a template (Figure 4). The snapback DNA synthesis structure provided direct evidence that G2L4 RT functions as dimer in a reaction akin to MMEJ but with both monomers activated at high concentrations of short single-stranded DNA substrate that would not reach the dimer interface. Modeling suggested that the conserved dimer interface of G2L4 RT enables it to bind to a 3’-DNA overhang on one side of a DSB site as a heterodimer consisting of a leading monomer activated by binding the 3’ end of the 3’-overhang as a primer and a trailing monomer that remains inactive, likely reflecting the situation at lower substrate concentrations *in vivo* (Figure 7). In this configuration, the NTE/RT0 loop of the active monomer is positioned to directly promote the annealing the 3’ end of the 3’-overhang on the left side of a DSB to a short microhomology at the 3’ end of the 3’-overhang on the right side of the break, while the NTE and the thumb domain of the trailing inactive monomer are positioned to further stabilize the 3’ overhang on left side of the DSB in an extended conformation that facilitates searching for a suitable microhomology in the 3’-overhang on the right side of the DSB that serves as a template. This model was supported by the finding that (i) mutations in interacting residues in the NTE portion of the G2L4 RT dimer interface that could impact the relative orientation of the NTE/RT0 loops of the two monomers virtually abolished the MMEJ activity of G2L4RT, while having much smaller effects on its primer extension and snapback DNA synthesis activities (Figure 6E and 6F); (ii) the NTE and the thumb domain of G2L4 RT contain conserved charged resides that are positioned in the trailing inactive monomer to stabilize the 3’-overhang on one side of the DSB in an extended conformation (Figures 7C-7E); and (iii) mutations in a subset of these residues preferentially inhibited MMEJ activity (Figure S7A and S7B).

Our structural and biochemical analyses revealed a series of conserved structural features that evolved to optimize the DSBR activity of G2L4 RT. These included the conserved I instead of A in the YxDD motif at the active site of G2L4 RT, which was found previously to favor strand annealing required for MMEJ over primer extension^20^. The snapback DNA synthesis structure showed that I238 forms a hydrophobic stacking interaction with the sugar pucker of the C+2 base, playing a pivotal role in securing the substrate within the active site (Figure 4G). This interaction maintains the nucleotide’s proper orientation, which is essential for accurate catalysis. I instead of A at the active site also favors the use of short annealed primers (2 to 5 nt) and strongly disfavors the use of longer annealed primers ≥10 nt^20^, consistent with the preference of the G2L4 RT MMEJ activity for shorter annealed microhomologies than GII RT^20^. Substitution of I with A at this position in G2L4 RT enabled the use of longer primers and annealed microhomologies^20^, possibly due to disruption of the stabilizing interaction of the Ile side chain with the sugar pucker of the incoming dNTP.

Other conserved structural features that optimize the DSBR activity of G2L4 RT included a longer NTE than that of GII RT with a smaller (5 nt) RT0 loop centered on three uniquely conserved Ser residues found in the snapback DNA synthesis structure to promote annealing of two 3’ nucleotides serving as a primer to two complementary residues in an upstream region serving as a DNA template (Figure 5D and E). This RT0 loop structure differs from that of GII RT, which forms a larger lid over positions -1 to +1 of a single-stranded region of the DNA template strand at the RT active site, as well as those for human LINE-1 and insect R2 element RTs that share the same “lid over the RT active site” morphology as GII RT (Figures S7E and S7F)^14,15,38^ All three of these RTs lack extended NTEs or a series of conserved serine residues that interact directly with a template strand, potentially a major difference between the basal DSBR activity of non-LTR-retroelement RTs and the optimized DSBR activity of G2L4 RTs.

Finally, the G2L4 RT structural features revealed in this study suggest new ways of optimizing non-LTR-retroelement RTs for genome engineering applications. These include: (i) modifications of RT3a that stabilize the free protein in an inactive configuration that prevents promiscuous reverse transcription of cellular RNAs until it encounters a physiological substrate; (ii) an extended NTE with an RT0 loop containing residues that interact directly with a single-stranded DNA overhang to form a stabilizing handle that favors strand annealing; (iii) the substitution of different residues at the x position Y/FxDD motif and its covarying partner in RT2a to modulate the balance between RT fidelity and strand annealing; and (iv) modifications of the outer surfaces of the NTE and thumb to form a dimer interface, whose stability could be tuned to localize two non-LTR-retroelement RT monomers to function together or separately for different genome engineering functions.

## Acknowledgements

This research was supported by NIH grant R01 CA281106 and R35 GM148356 to YJZ. and Welch Foundation grant F-1607 and NIH grant R35 GM136216 to AML. We thank Dr. Georg Mohr (University of Texas at Austin) for comments on the manuscript.

## AUTHOR CONTRIBUTIONS

SKP., MG., YJZ, and AML designed the studies and wrote the manuscript with help from JLS and SL. JLS performed protein purifications and crystallizations and solved the structure of G2L4 RT apoenzyme by X-ray crystallography. MG performed protein purifications and crystallization and solved the structure of snapback DNA substrate bound by G2L4 RT by X-ray crystallography with help from WK. SKP and SL performed protein purifications and biochemistry experiments.

## DECLARATION OF INTERESTS

The authors declare no competing interests.

## CONTACTS FOR REAGENT AND RESOURCE SHARING

Further information and requests for reagents may be directed to and will be fulfilled by the Lead Contact, Alan M. Lambowitz

## Methods

### Bacterial strains

*E. coli* HMS174 (DE3) (F^-^ recA1 hsdR (r_K12_^-^ m_K12_^+^) Rif ^R^ (DE3)) was purchased from Novagen and used for expressing proteins from recombinant plasmids. *E. coli* Rosetta 2 (F^-^ ompT hsdS_B_ (r_B_^-^ m_B_^-^) gal dcm pRARE2 (Cap^R^)) was purchased from Novagen and used for expression and purification of recombinant WT and mutant G2L4 RT proteins.

### Oligonucleotides

HPLC-purified oligonucleotides purchased from Integrated DNA Technologies (IDT) are listed in Table S1. For biochemical experiments, oligonucleotides were 5’-labeled with [γ-^32^P]-ATP (6,000 Ci/mmol; Revvity) by using T4 polynucleotide kinase (New England Biolabs) followed by clean-up with an Oligo Clean & Concentrator Kit (Zymo Research) according to the manufacturer’s protocols. Quantification of labeled oligonucleotides was performed by using a Qubit ssDNA assay kit according to the manufacturer’s protocol.

### Recombinant plasmids

pMal-c5X (New England Biolabs), which has a factor Xa cleavable maltose-binding protein (MBP) tag, an Amp^R^ marker, and isopropyl β-D-1-thiogalactopyranoside (IPTG)-inducible tac promoter was used to express pMal-G2L4 RT as described previously^20^. pMal-G2L4 RT mutants were constructed in the same expression plasmid by using a Q5 mutagenesis kit (New England Biolabs).

### Bioinformatics and amino acids sequence alignment for G2L4 RT and other RTs

The amino acid sequences of GII and G2L4 RTs were obtained from references 4 and 20, respectively. WebLogos^46^ for each RT were created by using 130 different G2L4 RTs and 500 different GsI-IIC (GII) RTs identified by BLASTP^47^ as having ≥50% sequence identity aligned using ClustalW via Jalview^48,49^. The NCBI accession codes for other RTs were sourced from Zimmerly and Wu (2015)^8^. Amino acids sequences of other RTs were obtained from NCIB or RCSB PDB (https://www.rcsb.org). Multiple sequence alignments of RTs were performed using T-Coffee through Jalview^48,50^.

### Protein purification

WT and mutant N-terminal MBP-tagged G2L4 RTs were purified as described^20^. pMal-c5x Amp^R^ WT and mutant G2L4 RT expression plasmids were transformed into *E. coli* Rosetta 2 Cap^R^ cells (Novagen). The transformants were plated on LB agar plates containing carbenicillin (100 µg/mL) and chloramphenicol (25 µg/mL) and incubated overnight at 37℃. The next day, a single colony was picked, inoculated into 20 mL of LB media containing carbenicillin (100 µg/mL) and chloramphenicol (25 µg/mL), and incubated with shaking at 220 rpm for 14-16 h at 37℃. The 20 mL-overnight grown cells were transferred into 1 L of LB media containing carbenicillin (100 µg/mL) and chloramphenicol (25 µg/mL) and incubated with shaking at 220 rpm at 37℃ until OD_600_ reached 0.8-1.0. WT and mutant G2L4 RTs were then induced by adding 100 µM IPTG and incubating at 100 rpm for 19-21 h at 18℃. After centrifugation in a 50-mL conical tube (Sarstedt) at 4,000 x g for 25 min, the cell pellets were transferred to another 50 mL- conical tube (Sarstedt) and resuspended in 45 mL of lysis buffer containing 20 mM Tris-HCl pH 7.5, 500 mM NaCl, 0.1% Triton X-100, 0.1% ß-mercaptoethanol, and 20% glycerol. The resuspended cells were then sonicated three times for 1 min at 80% amplitude using a Branson Sonifier 250 (Branson Ultrasonics), followed by centrifugation at 15,500 x g for 25 min. After collecting the supernatant, 0.04% polyethyleneimine (PEI; final concentration) was added, and the tube was inverted 2 or 3 times and then placed on ice for 10 min to precipitate nucleic acids. Precipitates were removed by centrifugation at 15,500 x g for 10 min, and the supernatant was filtered through a 0.45-mm pore size PES membrane (Thermo Fisher Scientific). The filtered supernatants were then loaded onto a 5-mL HiTrap MBP HP column (Cytiva) at flow rate of 5 mL/min using a AKTA^TM^ start FPLC (Cytiva), and the column was washed sequentially with 5 column volumes of buffer A (20 mM Tris-HCl pH 7.5, 100 mM NaCl, 0.1% ß-mercaptoethanol, and 10% glycerol) followed by 5 column volumes of washing buffer (20 mM Tris-HCl pH 7.5, 1.5 M NaCl, 0.1% Triton X-100, 0.1% ß-mercaptoethanol and 10% glycerol) and 5 column volumes of buffer A. Column-bound protein was eluted with 10 column volumes of elution buffer (20 mM Tris-HCl pH 7.5, 100 mM NaCl, 0.1% ß-mercaptoethanol, 10% glycerol, and 10 mM maltose). Column fractions containing the RT protein were identified by SDS-PAGE and staining with 0.25% Coomassie brilliant blue R (Sigma-Aldrich). The pooled fractions were loaded onto a 5-mL HiTrap Heparin HP column (Cytiva) at flow rate of 5 mL/min using a AK-TA^TM^ start FPLC (Cytiva). After washing the column with 5 column volumes of buffer A, bound proteins were eluted using 10 column volumes of an NaCl gradient between buffer A, which contains 100 mM NaCl, and buffer B, which contains 1.5 M NaCl (see above). Column fractions containing the RT protein were identified by SDS-PAGE and then pooled and concentrated using an Amicon® Ultra-15 (30k) concentrator (Millipore). The purified proteins were diluted in buffer containing 20 mM Tris-HCl pH 7.5, 50 mM NaCl and 50% glycerol to a final concentration of 8-10 mg/ml and stored at -80℃ until used.

### SDS- and native-polyacrylamide gel electrophoresis

For SDS-PAGE, 1 µg of WT or mutant G2L4 RTs were mixed with 5 µL of 4X sample buffer (200 mM Tris-HCl pH 6.8, 400 mM dithiothreitol (DTT), 8% SDS, 6 mM bromophenol blue, 40% glycerol) followed by addition of double-distilled H_2_O up to a volume of 20 µL and incubating at 95°C for 5 min. The samples were loaded on a NuPAGE 4-12% Bis-Tris gel with a 250 kDa Plus Prestained Protein Marker (Vazyme) in a parallel lane and run in 1X MES running buffer (Thermo Fisher Scientific) at 150 V for 1 h by using an XCell Surelock Electrophoresis Cell according to the manufacturer’s protocol. For the native gel, WT or mutant G2L4 RTs (1 µg/14 µL) was mixed with 3 µL of 6X native gel sample buffer (300 mM Tris-HCl pH 8.5, 60% glycerol) and 1 µL of 1% Coomassie Brilliant Blue G-250 solution (0.1 g Coomassie Brilliant Blue G-250 in 10 mL of 95% ethanol), followed by double-distilled H_2_O up to 18 µL. The samples were loaded on a 4-15% Criterion TGX Precast gel (Bio-Rad) and run in 1X Novex Tris-Glycine Native Running Buffer (Thermo Fisher Scientific) mixed with 0.1X NativePAGE Cathode Buffer Additive (Thermo Fisher Scientific) at 12 W for 1 h at 4℃ using a Criterion Cell (Bio-Rad) according to the manufacturer’s protocol.

### Factor Xa cleavage of the N-terminal MBP tag

To prepare G2L4 RT for crystallization, the N-terminal MBP tag was removed by incubating 50 µg/mL MBP-G2L4 RT with 1 µg/mL of Factor Xa (New England Biolabs) in 20 mM Tris-HCl pH 7.5, 50 mM NaCl, 2 mM CaCl_2_ at 4℃ overnight. Following cleavage, G2L4 RT was loaded onto a 5-mL HiTrap MBP HP column (Cytiva) at flow rate of 5 mL/min using a AKTA^TM^ start FPLC (Cytiva). The flow-through was collected and then loaded on a 5-mL HiTrap Heparin HP column (Cytiva) at flow rate of 5 mL/min, and the column was washed with 5 column volumes of buffer A. Bound proteins were eluted with a 10-column volume NaCl gradient using buffer A and buffer B (see above). Column fractions containing the RT protein were identified by SDS-PAGE, pooled, and protein concentration was measured by Bradford Protein assay kit (Bio-Rad) by using the manufacturer’s protocol. The pooled protein was concentrated to 8-9 mg/mL in crystallization buffer containing 5 mM Tris-HCl pH 7.5, 50 mM NaCl, 5 mM DTT and 10% glycerol using an Amicon Ultra-15 (30k) concentrator (Millipore Sigma) at 4℃ and stored at - 80℃ until used.

### G2L4 RT crystallization

For crystallization of G2L4 RT apoenzyme, a sitting drop mixture containing 1 µL of 10 mg/mL G2L4 RT without an MBP tag was incubated at 22℃ with an equal volume of crystallization buffer containing 0.1 M Bis-Tris pH 5.5, 0.2 M ammonium acetate and 25% polyethyleneglycol 3350 in an Intelli-Plate 96-2 Low Profile plate (Hampton Research). Crystals were observed after 5-8 days and harvested after 1 month. For co-crystallization, G2L4 RT without an MBP tag was prepared in crystallization buffer and mixed with a 15-nt snapback oligonucleotide (IDT; Table S1) at a 1:1.2 molar ratio of protein to ligand in a final volume of 100 µL. The mixture also included 20 mM MgCl₂, 1 mM MnCl₂, 2 mM dCTP (New England Biolabs), and 1 mM ddGTP (Millipore Sigma). After incubation at room temperature for 30 min, the mixture was set up in crystal trays. The final crystals were obtained in 0.2 M magnesium chloride hexahydrate, 0.1 M Bis-Tris pH 6.5, 25% polyethylene glycol 3350 (Hampton Research) at 25℃ after 5-8 days and were harvested after 16 days. Both apoenzyme G2L4 RT and co-crystal of G2L4 RT with a snap-back DNA oligonucleotide, dCTP and ddGTP were mounted by looping directly from the crystallization drop and plunging into liquid nitrogen for cryo-protection.

### G2L4 RT apoenzyme and snapback complex data collection, analysis, and structure determination

Diffraction data for G2L4 RT apoenzyme were collected on beamline 5.0.1 of the Advanced Light Source (ALS) at 100K from Lawrence Berkeley National Laboratory. Images were integrated using the XDS package^51^ and scaled with Aimless^52^. Initial structural determination was carried out using seleno-methionine single-wavelength anomalous diffraction (SAD) phasing on 720 degrees of data from one crystal that diffracted to 2.9 Å and showed a weak anomalous signal to 4 Å. Native crystal diffraction data were also collected in the same beamline (Table 1). ShelxC/D followed by Autosol from the Phenix package^53^ was used to generate a density-modified map that showed the three-helix bundle of the thumb and some clearly visible regions of the palm. Partial model building of the thumb and palm region was performed in the SAD density-modified map in Coot^54^ and refined in Refmac5^55^, and then used as a molecular replacement model in Phaser for the higher quality 2.6-Å native dataset. Further rounds of model building and refinement were carried out using a combination of Buster^56^ and Phenix, applying bulk solvent parameters, NCS restraints, TLS, and individual temperature factors. Diffraction data for G2L4 RT snapback complex were collected on beamline 5.0.2 of the Advanced Light Source (ALS) at 100K. X-ray diffraction data were processed to a resolution of 2.77 Å using HKL2000^57^ (Table 1). The molecular replacement solution for the complex structure was iteratively built using Coot with the Phenix refine package^58^. Final refinement was completed using Refmac ^55^. The quality of the final refined structure was evaluated by MolProbity^59^. Statistics for data collection and structure determination are shown in Table 1.

**Table 1.** Crystallographic Data Collection and Refinement Statistics.

| Table 1. Crystallographic Data Collection and Refinement Statistics |  |  |  |
| --- | --- | --- | --- |
|  | RT apoenzyme<br>(PDB: 9D5X) | RT snapback complex<br>(PDB: 9D5S) | RT(Se-SAD) |
| **Wavelength (Å)** | 0.9774 | 1.0000 | 0.9774 |
| **Resolution range (Å)** | 48.75-2.61 (2.70-2.61) | 47.30-2.77 (2.82-2.77) | 40.78-2.90 (3.00-2.90) |
| **Space group** | P2<sub>1</sub> | P2<sub>1</sub>2<sub>1</sub>2<sub>1</sub> | P2<sub>1</sub> |
| **Unit cell: a, b, c (Å)** | 75.836, 73.227, 82.861 | 65.398, 99.190, 157.276 | 75.077, 75.170, 83.012 |
| **Unit cell: $\alpha, \beta, \gamma$ (°)** | 90.00, 103.97, 90.00 | 90.00, 90.00, 90.00 | 90.00, 103.93, 90.00 |
| **Total reflections** | 182519 (17706) | 192211(191330) | 40010 (3973) |
| **Unique reflections** | 26981 (2684) | 26117 (2461) | 20027 (1988) |
| **Multiplicity** | 6.8 (6.6) | 7.2 (6.8) | 2.0 (2.0) |
| **Completeness (%)** | 99.77 (99.89) | 99.40 (99.30) | 99.74 (99.45) |
| **Mean I/sigma** | 21.11 (2.92) | 11.86 (1.29) | 42.65 (9.18) |
| **Wilson B- factor (Å<sup>2</sup>)** | 64.7 | 56.2 | 66.6 |
| **R-merge** | 0.056 (0.704) | 0.140 (0.836) | 0.013 (0.072) |
| **R-meas** | 0.061 (0.764) | 0.150 (0.904) | 0.019 (0.101) |
| **R-pim** | 0.023 (0.296) | 0.055 (0.339) | 0.013 (0.072) |
| **CC<sub>1/2</sub>** | 0.999 (0.862) | 0.991 (0.782) | 1.000 (0.994) |
| **CC\*** | 1.000 (0.962) | 0.998 (0.937) | 1.000 (0.999) |
| **Refinement** |  |  |  |
| **Reflections used in refinement** | 26935 (2683) | 26117 (2461) |  |
| **Reflections used for R-free** | 1356 (121) | 1960 (188) |  |
| **R-work** | 0.2167 (0.3823) | 0.2341 (0.3078) |  |
| **R-free** | 0.2430 (0.4409) | 0.2731 (0.3436) |  |
| **CC (work)** | 0.971 (0.768) | 0.925 (0.790) |  |
| **CC (free)** | 0.964 (0.728) | 0.924 (0.718) |  |
| **Number of non-hydrogen atoms** | 6236 | 7010 |  |
| **Macromolecules** | 6184 | 6952 |  |
| **Protein residues** | 782 | 816 |  |
| **RMS (bonds) (Å)** | 0.002 | 0.011 |  |
| **RMS (angles) (°)** | 0.43 | 1.53 |  |
| **Ramachandran favored (%)** | 98.05 | 97.78 |  |
| **Ramachandran allowed (%)** | 1.69 | 1.97 |  |
| **Ramachandran outliers (%)** | 0.26 | 0.25 |  |
| **Average B- factor (Å<sup>2</sup>)** | 79.0 | 63.1 |  |
| **Macromolecules** | 79.1 | 63.0 |  |
| **Ligands** | 67.8 | 50.0 |  |
| **Molprobity score** | 1.09/100<sup>th</sup> percentile | 1.74/100<sup>th</sup> percentile |  |
| Statistics for the highest-resolution shell are shown in parentheses.<br>Se-SAD, seleno-methionine single-wavelength anomalous diffraction.<br>CC<sub>1/2</sub> is the Pearson correlation coefficient for a random half of the data, the two numbers represent the lowest and highest resolution shell, respectively.<br>MolProbity score is calculated by combining clashscore with rotamer and Ramachandran percentage and scaled based on X-ray resolution. The percentage is calculated with 100th percentile as the best and 0th percentile as the worst among structures of comparable resolution.<br>The outlier for both apoenzyme and complex structure is I238, a crucial active site residue. The density is clear and the unique phi/psi angle might be related to its role in activity. |  |  |  |

### Structural figures and map visualization

All structural figures were created using USCF ChimeraX (version 1.8)^60^. The Fo-Fc map was calculated using Coot^54^ and visualized using UCSF ChimeraX (version 1.8)^60^. The map was contoured at 3 σ to highlight significant differences between the observed and modeled electron density. The 2Fo-Fc map was generated using Coot^54^ and visualized using UCSF ChimeraX (version 1.8)^60^. The map was contoured at 1.5 σ to visualize the electron density, aiding in model refinement and validation. Positive densities confirmed the presence of modeled atoms, while absent or poorly contoured densities suggested model inaccuracies.

### Differential Scanning Fluorimetry

Roche Light Cycler® 480II was used for Differential Scanning Fluorimetry (DSF) assays. SYPRO Orange dye (Thermo Fisher Scientific, 5,000x in DMSO) was used for fluorescence emission. The reaction was prepared in a total volume of 20 μL with a final protein concentration of 5 μM. The dilution buffer used was identical to the gel filtration buffer, consisting of 20 mM Tris-HCl pH 7.5, 50 mM NaCl, 0.1% % ß-mercaptoethanol, 10% glycerol. SYPRO Orange dye (5000x stock solution) added to the reaction mix at a final concentration of 1x. The mixture was placed in a 96-well plate designed for a real-time PCR instrument (Life Technologies), thoroughly mixed, incubated for 10 min at room temperature, and then centrifuged at 2,000 × g for 5 min. The assay plate was placed into a Roche Light Cycler® 480II to obtain a pre-programmed experimental Melting Curve with target temperature set for 95℃, retention time set at continuous, and ramp rate at 0.04℃/second. The software analysis function was used to calculate melting temperature (Tm) for each sample.

### Size-exclusion chromatography

Wild-type and mutant G2L4 RTs with or without the N-terminal MBP tag were purified as described above. The oligomeric state of the proteins was evaluated using HiLoad 16/600 Superdex 200 (Cytiva) and Superdex 75 prep grade columns (Cytiva) for proteins with or without the MBP tag, respectively. Concentrated protein samples were injected onto the columns using a Bio-Rad chromatography system. Elution was carried out at a flow rate of 0.8 mL/min with a running buffer comprised of 20 mM Tris-HCl pH 7.5, 50 mM NaCl, 0.1% β-mercaptoethanol, and 10% glycerol at 4°C. UV absorbance at 280 nm was monitored with a Bio-Rad detector to track the elution profiles. Molecular weights were calculated based on the calibration curve for elution volumes versus the logarithm of the molecular weight of Gel Filtration Standard (Bio-Rad). Fractions were collected for further analysis, and the molecular weights of the proteins in solution were determined using a previously established calibration curve to assess their oligomeric state. Following size-exclusion chromatography, proteins were concentrated to 8-9 mg/mL in a crystallization buffer containing 5 mM Tris-HCl pH 7.5, 50 mM NaCl, and 10% glycerol for subsequent crystallization experiments.

### Primer extension assays

Biochemical assays were done using G2L4 RTs stabilized with an N-terminal MBP-tag. Primer extension assays were done using a 50-nt DNA template oligonucleotide ending with an inverted 3’ dT that blocks use for snapback DNA synthesis or terminal transferase activity (IDT; Table S1). The DNA template (1 µM) was pre-annealed to 200 µM of a 5-nt DNA primer in 100 µL of TE (10 mM Tris-HCl pH 7.5, 1 mM EDTA) by heating to 95°C for 3 min followed by cooling to 25°C at 0.1°C /min in a T100 thermal cycler (Bio-Rad). The assays were performed in 80 µL of reaction medium containing 500 nM WT or mutant G2L4 RTs, 250 nM template-primer complex, 20 mM Tris-HCl pH 7.5, 20 mM NaCl, and 10 mM MgCl_2_ with or without 1 mM MnCl_2_. After pre-incubating the RT with the annealed template-primer substrate for 30 min at room temperature, the reactions were initiated by adding 1 mM dNTPs (1 mM each of dATP, dCTP, dGTP and dTTP) plus 1 µCi [α-^32^P]-dTTP (3,000 Ci/mmol; Revvity) and incubated at 37°C for times up to 180 min. For time courses, 10-µL aliquots were taken at each time point and quenched by adding 2 µL of 6X stop solution (25 mM EDTA, 0.5 U/µL Proteinase K (NEB)) and incubating for 15 min at 37°C. The samples were then mixed with an equal volume of 2X RNA loading dye (95% formamide, 0.02% SDS, 0.02% bromophenol blue, 0.01% xylene cyanol, 1 mM EDTA) and analyzed by electrophoresis in a 20% polyacrylamide gel with TBE (89 mM Tris, 89 mM Borate and 2 mM EDTA) buffer against 5’-labeled synthesized DNA markers. The gel was dried and scanned using a phosphorimager (Typhoon FLA 9500; GE Healthcare) and processed with ImageJ^61^. The amount of labeled dTTP incorporated into the product was quantified using ImageQuant TL 10.2. To account for differences in primer extension product sizes, the label amount was normalized by multiplying the dTTP concentration (1 mM) and dividing by the number of T residues (22 bases) per extension product. This provided the concentration of extended product, which was then plotted relative to the template concentration (250 nM). Time course data were fit to a first-order rate equation using Prism 10.0 to obtain k_obs_ and amplitude values. For slow reactions without a clear end point, the reaction amplitude was fit to match average end point of parallel reactions that reached clear end points. Amplitude values obtained in this way are indicated in parenthesis in tables next to the plots.

### Snapback DNA synthesis assays

The substrate for snapback replication assays was a 50-nt DNA oligonucleotides with an unblocked 3’ OH end (Table S1). The assays were done as time courses by pre-incubating 10 nM 5’-^32^P -labeled 50-nt DNA oligonucleotide with 500 nM enzyme in 80 µL of the reaction medium containing 20 mM Tris-HCl pH 7.5, 20 mM NaCl, 10 mM MgCl_2_ and with or without 1 mM MnCl_2_ for 30 min at room temperature. Reactions were initiated by adding 1 mM dNTPs (an equimolar mix of dATP, dCTP, dGTP, and dTTP). 10-µL aliquots were taken at each time point up to 180 min, quenched with 2 µL of 6X stop solution, and analyzed by electrophoresis in a non-denaturing 12% polyacrylamide gel with dsDNA size markers, a 5’-labeled Low Molecular Weight DNA Ladder (New England Biolabs) with TBE (89 mM Tris, 89 mM Borate and 2 mM EDTA) buffer. The gel was dried, scanned with a phosphorimager (Typhoon FLA 9500; GE Healthcare), and processed with ImageJ. Products were quantified with ImageQuant TL 10.2, and data were analyzed as described above for primer extension assays.

### Microhomology-mediated end-joining assays

MMEJ assay was performed as described^20^ with minor modifications. The MMEJ assays were done using a partially double-stranded DNA consisting of a 5’-labeled 53-nt oligonucleotide (D1) ending with a 4-nt microhomology annealed to an unlabeled complementary 39-nt DNA oligonucleotide with a 3’ inverted dT blocking group (D2) leaving a 15-nt single-stranded 3’ overhang. To anneal complementary strands in the MMEJ substrate, 10 µM D1 and 20 µM D2 oligonucleotides in 100 µL TE buffer were heated to 95°C for 3 min and cooled at 0.1°C/min to 25°C in a T100 thermal cycler (Bio-Rad). In Figures, the annealed products are denoted D1/D2 and D1’/D2’. Reactions were done using 250 nM of the annealed substrate preincubated with 500 nM WT or mutant G2L4 RT in the reaction medium containing 20 mM Tris-HCl pH 7.5, 20 mM NaCl, and 10 mM MgCl_2_ with or without 1 mM MnCl_2_ for 30 min at room temperature. The reactions were initiated by adding 1 mM dNTPs (an equimolar mix of dATP, dCTP, dGTP, and dTTP), incubated at 37°C for up to 180 min, and quenched as described for primer extension assays. Products were analyzed by electrophoresis on a non-denaturing 12% polyacrylamide gel against double-stranded DNA size markers (5’-labeled Low Molecular Weight DNA Ladder; New England Biolabs) with TBE (89 mM Tris, 89 mM Borate and 2 mM EDTA) buffer. The gel was scanned with a phosphorimager (Typhoon FLA 9500; GE Healthcare) and processed using ImageJ. Products were quantified with ImageQuant TL 10.2, and data were analyzed as described for primer extension assays.

### Terminal transferase assays

Terminal transferase assays were done using the same 50-nt DNA oligonucleotides used as templates in primer extension assays but 5’-^32^P-labeled and without a 3’-blocking group (Table S1). The reactions were done as 30-min time courses in 80 µL of reaction medium. WT or mutant G2L4 RT (500 nM) was pre-incubated with 5’-labeled oligonucleotide substrate (10 nM) in reaction medium containing 20 mM NaCl, 10 mM MgCl_2_, and 20 mM Tris-HCl pH 7.5 with or without 1 mM MnCl_2_ for 30 min at room temperature. The reaction was initiated by adding 1 mM of a single dNTP (dATP, dCTP, dGTP, or dTTP) and incubating at 37°C for times indicated for individual experiments, then quenched as described for primer extension assays. Products were analyzed on a denaturing 6% polyacrylamide gel with a 5’-labeled RiboRuler Low Range RNA Ladder (Thermo Fisher Scientific) as size markers with TBE (89 mM Tris, 89 mM Borate and 2 mM EDTA) buffer. The gel was dried and scanned with a phosphorimager (Typhoon FLA 9500; GE Healthcare), and processed using ImageJ. Quantification was done with ImageQuant TL 10.2, and data were analyzed as described for primer extension assays.

### Modeling of MMEJ substrate for hypothetical G2L4 RT DSBR mechanism

Unwound DNA from the superfamily 2 helicase Hel308 complex (PDB ID: 2P6R)^62^ was used as a structural basis to model the MMEJ substrate in the G2L4 RT-mediated DSBR pathway. The partially unwound DNA in this structure consisted of a 5 to 3′ 25-mer strand and a 3′ to 5′ 15-mer strand. Using Coot, the last two unpaired nucleotides of the 15-mer strand were trimmed for our model to generate a partial duplex with a 12-nt single-stranded 3’-overhang. To prepare the MMEJ substrate model, one copy was modified by shortening the 3′ single-stranded overhang to 10 nucleotides and changing the last five nucleotides to AACCG. This modified strand was then superimposed onto the primer strand of a snapback substrate using the Coots LSQ superimposition function. Another copy of the duplex structure was altered by generating four additional nucleotides along the 3′ single-stranded overhang and mutating the last six nucleotides to GCGGTT. This modified strand was superimposed onto the template strand of a snapback substrate using the same LSQ superimposition function. Following alignment, overlapping nucleotides at the 3′ ends of the modified strands were removed to eliminate redundancy. The three molecules were then merged, and Coot’s Real Space Refinement function was applied to connect the original substrate and the additional partial duplex, resulting in the generation of two new chains: one 29 nt in length, corresponding to the modified template strand, and the other 23 nt in length, corresponding to the modified primer strand. The resulting structural model was subsequently prepared using the Protein Preparation Workflow in Maestro (Schrödinger Suite version 2023-1)^63^ with default parameters. This workflow facilitated energy minimization to refine the geometry of the model.

### Data Resources

The X-ray crystallography structures of G2L4 RT apoenzyme and G2L4 RT bound to a snapback DNA substrate have been deposited in the Protein Data Bank under accession number as PDB: 9D5X (apoenzyme) and 9D5S (snapback substrate bound), respectively.

## Acknowledgements

We thank the Electron Microscopy Facility of the National Institute of Neurological Disorders and Stroke, the Central Microscopy Facility of the Marine Biological Laboratory, Christine A. Winters (NINDS) and Katherine Hammar (MBL) for sample preparations and technical support. We thank Professor Adriano Senatore, University of Toronto, for valuable comments on the manuscript. This work was supported by the Intramural Research Program of the National Institute of Neurological Disorders and Stroke, National Institutes of Health, National Institutes of Health, Bethesda, MD, USA.

---

**Figure S1.** Characteristics and comparisons of G2L4 and GII RTs related to Figure 1. (A) Superdex 200 size-exclusion chromatography profiles of purified MBP-tagged G2L4 RT (black) and GII RT (red). Vertical dashed lines indicate peak elution volumes for MBP-G2L4 and MBP-GII RTs. Molecular weights of the proteins based on the peak elution volumes were calculated using a calibration curve for log_10_ of the molecular weights (MW) of Superdex 200 protein standards shown to the right. (B) Superimposition of the two monomer subunits of G2L4 RT in the X-ray crystal structure of apoenzyme dimers. Regions of the monomer A are color-coded as in Figure 1A and monomer B is colored gray. (C) Sequence alignment of G2L4 and GII RT done by ClusterW via Jalview^48,49^. WebLogos are based on 130 G2L4 RTs and 500 GII RTs searched by BLASTP for ≥50% Identity of amino acids sequence and aligned by ClustalW. Sequence motifs found in all RTs (RT1-7) and NTE/RT0 loop, RT2a, and RT3a found in non-LTR-retroelement RTs are delineated above and highlighted in blue boxes. The YxDD motif at the RT active site is boxed in green. Red boxes indicate insertions in G2L4 RT relative to GII RT. S412 of G2L4 RT is derived from the expression plasmid. Amino acid residues in α-helical (H), random coil (C), and b-sheet (E) regions in the indicated crystal structures are indicated below the amino acid sequence. α1 through α5 are thumb domain α-helices in G2L4 RT labeled in Figure 1B. The last 2 amino acids of G2L4 RT were not visible in the substrate bound active structure.

**Figure S2.** Sequence alignments and structural analysis of adjacent RT3a and RT4/5 regions of G2L4 and other RTs related to Figure 2. (A) Multiple sequence alignment of the RT3a knot region of G2L4 RT (top) with those of other RTs (NCBI accession numbers or PDB codes to the right of the protein names) performed by T-Coffee via Jalview^48,50^. Other RTs include group IIC, IIA, and IIB intron RTs; bacterial chromosomally encoded group II intron-like 1, 2, 3, and 5 RTs; non-LTR-retrotransposon RTs; other chromosomally encoded bacterial RTs, including RVT, which is also found in some eukaryotes; and HIV-1 RT. The G2L4 RT3a knot region is highlighted in a purple box, with the 9-aa insertion within the G2L4 RT3a knot highlighted in a black box. Amino acid residues in boxes are colored-coded by their hydropathic character as charged (blue), polar (purple) or hydrophobic (red)^64^. The numbers at the top indicate the positions of amino acids in the G2L4 RT sequence. Numbers in parentheses in gaps in the alignments for some RTs indicate the number of additional amino acids not shown in the figure. (B) Amino acids sequence alignments and WebLogos comparing the RT3a, RT4, and RT5 regions of G2L4 RTs to *Roseburia intestinalis* (R.i) and *Eubacterium rectale* (E.r.) group IIC intron RTs, which lack an RT3a knot. The RT3a knot and its 9-aa insertion in G2L4 RT are highlighted in the alignment to illustrate their evolutionary conserved amino acid sequence in 130 different G2L4 RTs. Amino acids in α-helical (H), random coil (C), and β-sheet (E) regions in crystal structures are indicated below the amino acid sequence. (C) Superimposition of crystal structures of the fingers and palm regions of the R.i. (PDB:5HHJ; left) and E.r. (PDB:5HHL; right) RTs^31^ colored as in Figure 1B with those of G2L4 RT (white) showing differences in the interaction of RT3a with the RT active site. The arrows point to the partially hidden PQG motifs (yellow) at the beginning of RT4 of the R.i. and E.r. RTs.

**Figure S3.** SDS-polyacrylamide gel electrophoresis of WT and RT3a mutant G2L4 RTs and further biochemical assays related to Figure 3. (A) Coomassie blue-stained NuPAGE 4-12% Bis-Tris gel of WT and RT3a mutant G2L4 RTs used in biochemical assay. Proteins were expressed with a N-terminal MBP tag and purified as described in Methods. The numbers to the left of the gel indicate molecular weights of a Color Prestained Broad Range (10-250 kDa) protein ladder (New England Biolabs). (B) Terminal transferase assay time courses with a 5’-^32^P-labeled 50-nt DNA substrate without a 3’-blocking group. Reactions were initiated by adding 1 mM of a single dNTP (dATP, dCTP, dGTP, and dTTP) and incubated at 37°C for times up to 30 min in reaction medium containing 10 mM Mg^2+^ in the absence (top panels) or presence (bottom panels) of 1 mM Mn^2+^. The numbers to the left of the gel indicate the positions of 5’-labeled RiboRuler Low Range RNA Ladder size markers run in a parallel lane. The plots to the right of the gel show the average value and variance for two repeats of the experiment. Tables to the right of the plots show the rate constants (k_obs_) and amplitudes (Ampl.) of labeled products >50 nt obtained by fitting to a first-order rate equation. Ampl. values in parentheses represent fixed amplitudes for reactions that did not reach an end point based on the average Ampl. value for those that reached a clear end point during the experiment. The red asterisk in the schematic at the bottom indicates a 5’-^32^P-label.

**Figure S4.** 2Fo-Fc map of G2L4 RT complex with the bound snapback DNA substate and incoming dNTP related to Figure 4. (A) The 2Fo-Fc map was calculated by doubling the observed electron density (Fo) and subtracting the calculated electron density (Fc) to provide an estimate of the true electron density of the active site of G2L4 RT with bound substrate. Positive density of the electron cloud is shown as mesh and colored in blue contoured at a level of 1.5 sigma (σ). Nucleotides are numbered and colored as in Figure 4A. (B) Superimposition of active conformations of G2L4 RT and GII RT. Different regions of G2L4 RT are color-coded as in Figure 1, and GII RT is colored gray. (C) Superimposition of G2L4 RT apoenzyme and snapback DNA synthesis structure focusing on the AQG motif in the snapback DNA complex (yellow) and apoenzyme (gray). The pink sphere is a divalent metal ion in the snapback DNA complex structure. (D) Close-up view of the active site region of GII RT^4^ highlighting key residues, including D223 (blue) and L139 involved in coordination of Mg^2+^ (green sphere) and binding of the incoming dATP (green base, orange triphosphates) by D223 (blue) and R75 (salmon). The DNA template (violet) and primer (cyan) are depicted as sticks.

**Figure S5.** Sequence alignments of the NTE/RT0 loop of G2L4 RT compared to other RTs and terminal transferase assays of WT and RT0 loop mutant G2L4 RTs related to Figure 5. (A) Multiple sequence alignments of the NTE/RT0 loop region of G2L4 RT (top) with those of other RTs (NCBI accession numbers or PDB codes to the right of protein names) performed by T-Coffee via Jalview^48,50^. The color scheme of amino acids boxes is based on their Chou-Fasman two turn propensity (red, highest turn propensity; cyan, lowest turn propensity)^65^. The numbers at the top indicate the positions of the G2L4 RT sequence. The asterisks indicate conserved serine residues in G2L4 RTs. (B) Terminal transferase assays of MBP-tagged G2L4 wild-type and S3/G3 mutant RTs in the presence of 10 mM Mg^2+^ (left assays) or 10 mM Mg^+2^ + 1 mM Mn^2+^ (right assays) using a 5’- labeled 50-nt DNA substrate, as described in Figures S3B. Reactions were initiated by adding 1 mM of a single dNTP (dATP, dCTP, dGTP, and dTTP) and incubated at 37°C for times up to 30 min. The numbers to the left of the gels indicate the positions of 5’-labeled RiboRuler Low Range RNA Ladder size markers run in a parallel lane. The plots show the average values and variance for two repeats of the experiment. The Tables to the right of the plots indicate rate constants (k_obs_) and amplitudes (Ampl.) for synthesized DNA products by WT and S3/G3 RT0 loop mutant G2L4 RTs with a curve fit to a first-order rate equation. Ampl. values in parentheses represent fixed amplitudes for reactions that did not reach an end point based on the average Ampl. value for those that reached a clear end point during the experiment. The red asterisk in the schematic at the bottom indicates a 5’-^32^P-label.

**Figure S6.** Profiling of G2L4 RT dimer interface mutants related to Figure 6. (A) Size-exclusion chromatography of WT G2L4 RTs with a cleaved MBP tag (left panel) and a G2L4 RT C-terminal deletion mutant (ι1401-411 amino acids; right panel). The molecular weights in parenthesis were calculated based on the elution volume of the peak (dashed line) relative to the protein standard calibration graph of Figure S1A. (B) Differential Scanning Fluorimetry (DSF) assays. Left panel, G2L4 RT with (black) or without (red) an MBP tag; middle panel, C-terminal deletion mutant of G2L4 RT (ι1401-411 amino acids, green) versus WT G2L4 RT (black); right panel, dimer interface NTE mutant (R12/Y15/R19A; red); Thumb (T) domain mutant (R397/R398/R404A; blue); and NTE+T mutant (R12/Y15/R398A; yellow). The plots show the derivative of fluorescence intensity as a function of temperature, highlighting transitions or melting points indicative of structural changes or stability shifts. (C) Coomassie blue-stained non-denaturing gel for WT and mutant G2L4 RTs with an N-terminal MBP tag. Dimer and monomer bands are labeled to the right. The fractions of monomer bands indicated below the gel were calculated by using ImageQuant TL 10.2 software. (D) Terminal transferase assays with or without Mn^2+^ of MBP-tagged G2L4 RT dimer interface mutants using a 5’-labeled 50-nt DNA substrate, as described in Figure S3B.

**Figure S7.** G2L4 RT biochemical assay and models supporting the suggested G2L4 RT MMEJ mechanism and RT0 loop interactions for human LINE-1 and insect R2 non-LTR-retroelements related to Figure 7. (A and B) Biochemical assays of G2L4 RT NTE mutants in reaction media containing 10 mM Mg^2+^ (panel A) or 10 mM Mg^2+^ plus 1 mM Mn^2+^ (panel B). Left, primer extension assays; right, MMEJ assays. The plots show the average values and variance for two repeats of the experiment. Ampl. values in parentheses represent fixed amplitudes for reactions that did not reach an end point based on the average Ampl. value for those that reached a clear end point during the experiment. (D) Model of a G2L4 RT dimer with two active monomers bound to a 5-bp annealed microhomology between the single-stranded 3’ overhangs from the left side (cyan) and right side (violet) of the DSB. A dashed red circle highlights a gap between single-stranded region of the MMEJ substrate and the trailing active monomer. (E) Model of a G2L4 RT dimer with an active and inactive monomers bound to a 7-bp annealed microhomology between the same single-strand 3’ overhangs as panel C. A steric clash is highlighted in a dashed red circle. (F) Wide-angle views of the RT0 region of LINE-1 RT (PDB: 8C8J)^15^ with bound RNA template/DNA primer substrate and dTTP. The RNA template (violet), DNA primer (cyan), and bound dTTP (light blue/orange) are depicted as sticks. Tower (light yellow) and Wrist (pink) are additional regions of LINE-1 RT. (G) Wide-angle views of the RT0 region of *Bombyx mori* R2 element RT (PDB:8GH6)^38^ modeled with the same DNA temple/DNA primer substrate as G2L4 RT. The DNA template (violet), DNA primer (cyan), and bound dTTP (blue base) and dCTP (yellow base) are depicted as sticks.

## Notes

### Competing Interest Statement

The authors have declared no competing interest.

