## Supplementary figures and table for "Structural basis for the evolution of a domesticated group II intron-like reverse transcriptase to function in host cell DNA repair"

### Figure S1

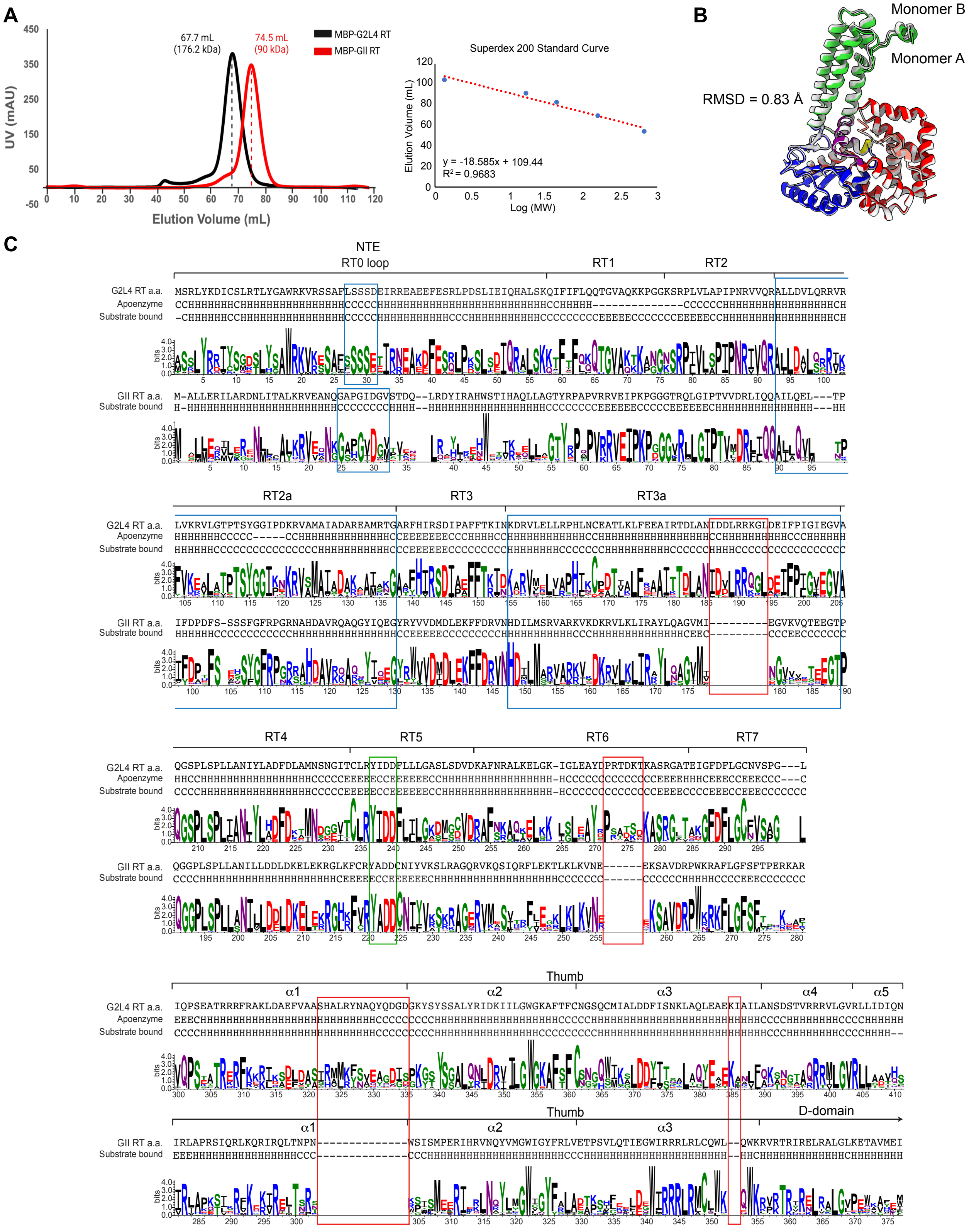
A
MBP-G2L4 RT
MBP-GII RT
67.7 mL
(176.2 kDa)
74.5 mL
(90 kDa)
UV (mAU)
Elution Volume (mL)
Superdex 200 Standard Curve
Elution Volume (mL)
Log (MW)
y = -18.585x + 109.44
R² = 0.9683
B
Monomer B
Monomer A
RMSD = 0.83 Å
C
NTE
RT0 loop
RT1
RT2
RT2a
RT3
RT3a
RT4
RT5
RT6
RT7
Thumb
α1
α2
α3
α4
α5
D-domain
G2L4 RT a.a.
Apoenzyme
Substrate bound
GII RT a.a.
bits

**Figure S1. Characteristics and comparisons of G2L4 and GII RTs related to Figure 1**

(A) Superdex 200 size-exclusion chromatography profiles of purified MBP-tagged G2L4 RT (black) and GII RT (red). Vertical dashed lines indicate peak elution volumes for MBP-G2L4 and MBP-GII RTs. Molecular weights of the proteins based on the peak elution volumes were calculated using a calibration curve for $\log_{10}$ of the molecular weights (MW) of Superdex 200 protein standards shown to the right.

(B) Superimposition of the two monomer subunits of G2L4 RT in the X-ray crystal structure of apoenzyme dimers. Regions of the monomer A are color-coded as in Figure 1A and monomer B is colored gray.

(C) Sequence alignment of G2L4 and GII RT done by ClusterW via Jalview[48,49]. WebLogos are based on 130 G2L4 RTs and 500 GII RTs searched by BLASTP for ≥50% Identity of amino acids sequence and aligned by ClustalW. Sequence motifs found in all RTs (RT1-7) and NTE/RT0 loop, RT2a, and RT3a found in non-LTR-retroelement RTs are delineated above and highlighted in blue boxes. The YxDD motif at the RT active site is boxed in green. Red boxes indicate insertions in G2L4 RT relative to GII RT. S412 of G2L4 RT is derived from the expression plasmid. Amino acid residues in α-helical (H), random coil (C), and b-sheet (E) regions in the indicated

crystal structures are indicated below the amino acid sequence. α1 through α5 are thumb domain α-helices in G2L4 RT labeled in Figure 1B. The last 2 amino acids of G2L4 RT were not visible in the substrate bound active structure.

#### Figure S2

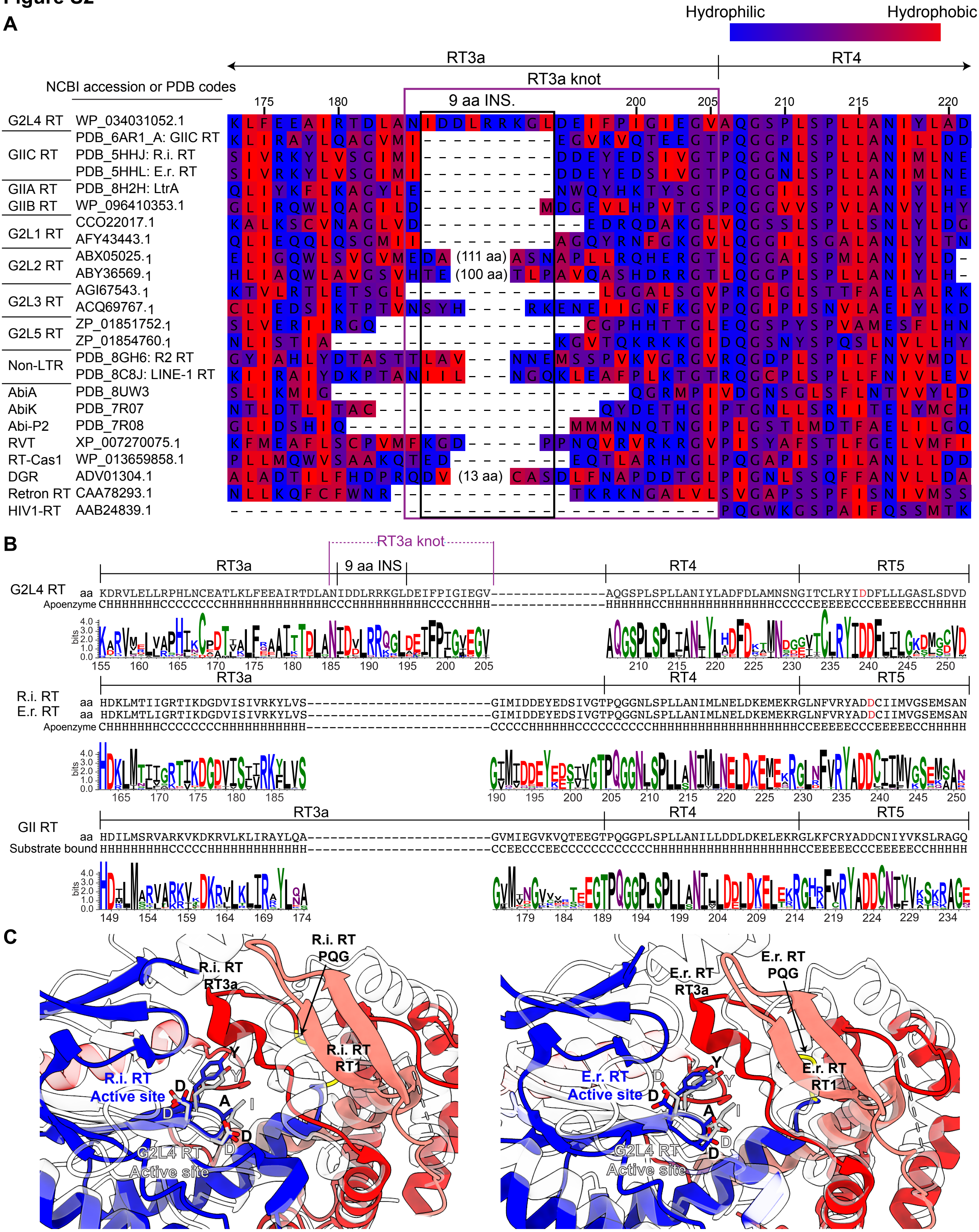
A
Hydrophilic
Hydrophobic
RT3a
RT4
RT3a knot
9 aa INS.
NCBI accession or PDB codes
G2L4 RT WP_034031052.1
GIIC RT PDB_6AR1_A: GIIC RT
PDB_5HHJ: R.i. RT
PDB_5HHL: E.r. RT
GIIA RT PDB_8H2H: LtrA
GIIB RT WP_096410353.1
G2L1 RT CCO22017.1
AFY43443.1
G2L2 RT ABX05025.1
ABY36569.1
G2L3 RT AGI67543.1
ACQ69767.1
G2L5 RT ZP_01851752.1
ZP_01854760.1
Non-LTR PDB_8GH6: R2 RT
PDB_8C8J: LINE-1 RT
AbiA PDB_8UW3
AbiK PDB_7R07
Abi-P2 PDB_7R08
RVT XP_007270075.1
RT-Cas1 WP_013659858.1
DGR ADV01304.1
Retron RT CAA78293.1
HIV1-RT AAB24839.1
(111 aa)
(100 aa)
(13 aa)
B
RT3a knot
RT3a
9 aa INS
RT4
RT5
G2L4 RT aa KDRVLELLRPHLNCEATLKLFEEAIRTDLANIDDLRRKGLDEIFPIGIEGV---------------AQGSPLSPLLANIYLADFDLAMNSNGITCLRYIDDFLLLGASLSDVD
Apoenzyme CHHHHHHHCCCCCCCCHHHHHHHHHHHHHHHCCHHHHHHHHHHCCCCHHHH---------------HHHHHHCCHHHHHHHHHHHHHHHHCCCCEEEEECCCEEEEEECCHHHHH
R.i. RT aa HDKLMTIIGRTIKDGDVISIVRKYLVS------------------------GIMIDDEYEDSIVGTPQGGNLSPLLANIMLNELDKEMEKRGLNFVRYADDCIIMVGSEMSAN
E.r. RT aa HDKLMTLIGRTIKDGDVISIVRKYLVS------------------------GIMIDDEYEDSIVGTPQGGNLSPLLANIMLNELDKEMEKRGLNFVRYADDCIIMVGSEMSAN
Apoenzyme CHHHHHHHCCCCCCCCHHHHHHHHHHH------------------------CCCCCHHHHHHCCCCCCCCCHHHHHHHHHHHHHHHHHHHHCCEEEEECCCEEEEEECCHHHHH
GII RT aa HDILMSRVARKVKDKRVLKLIRAYLQA------------------------GVMIEGVKVQTEEGTPQGGPLSPLLANILLDDLDKELEKRGLKFCRYADDCNIYVKSLRAGQ
Substrate bound HHHHHHHHHCCCCCHHHHHHHHHHHHH------------------------CCEECCCEECCCCCCCCCCCCHHHHHHHHHHHHHHHHHHHHCCEEEEECCCEEEEEECCHHHHH
bits
C
R.i. RT PQG
R.i. RT RT3a
R.i. RT RT1
R.i. RT Active site
G2L4 RT Active site
E.r. RT PQG
E.r. RT RT3a
E.r. RT RT1
E.r. RT Active site

**Figure S2. Sequence alignments and structural analysis of adjacent RT3a and RT4/5 regions of G2L4 and other RTs related to Figure 2**

(A) Multiple sequence alignment of the RT3a knot region of G2L4 RT (top) with those of other RTs (NCBI accession numbers or PDB codes to the right of the protein names) performed by T-Coffee via Jalview[48,50]. Other RTs include group IIC, IIA, and IIB intron RTs; bacterial chromosomally encoded group II intron-like 1, 2, 3, and 5 RTs; non-LTR-retrotransposon RTs; other chromosomally encoded bacterial RTs, including RVT, which is also found in some eukaryotes; and HIV-1 RT. The G2L4 RT3a knot region is highlighted in a purple box, with the 9-aa insertion within the G2L4 RT3a knot highlighted in a black box. Amino acid residues in boxes are colored-coded by their hydropathic character as charged (blue), polar (purple) or hydrophobic (red)[64]. The numbers at the top indicate the positions of amino acids in the G2L4 RT sequence. Numbers in parentheses in gaps in the alignments for some RTs indicate the number of additional amino acids not shown in the figure.

(B) Amino acids sequence alignments and WebLogos comparing the RT3a, RT4, and RT5 regions of G2L4 RTs to *Roseburia intestinalis* (R.i) and *Eubacterium rectale* (E.r.) group IIC intron RTs, which lack an RT3a knot. The RT3a knot and its 9-aa insertion in G2L4 RT are highlighted in the alignment to illustrate their evolutionary conserved amino acid sequence in 130 different G2L4 RTs. Amino acids in $\alpha$-helical (H), random coil (C), and $\beta$-sheet (E) regions in crystal structures are indicated below the amino acid sequence.

(C) Superimposition of crystal structures of the fingers and palm regions of the R.i. (PDB:5HHJ; left) and E.r. (PDB:5HHL; right) RTs[31] colored as in Figure 1B with those of G2L4 RT (white) showing differences in the interaction of RT3a with the RT active site. The arrows point to the partially hidden PQG motifs (yellow) at the beginning of RT4 of the R.i. and E.r. RTs.

**Figure S3**

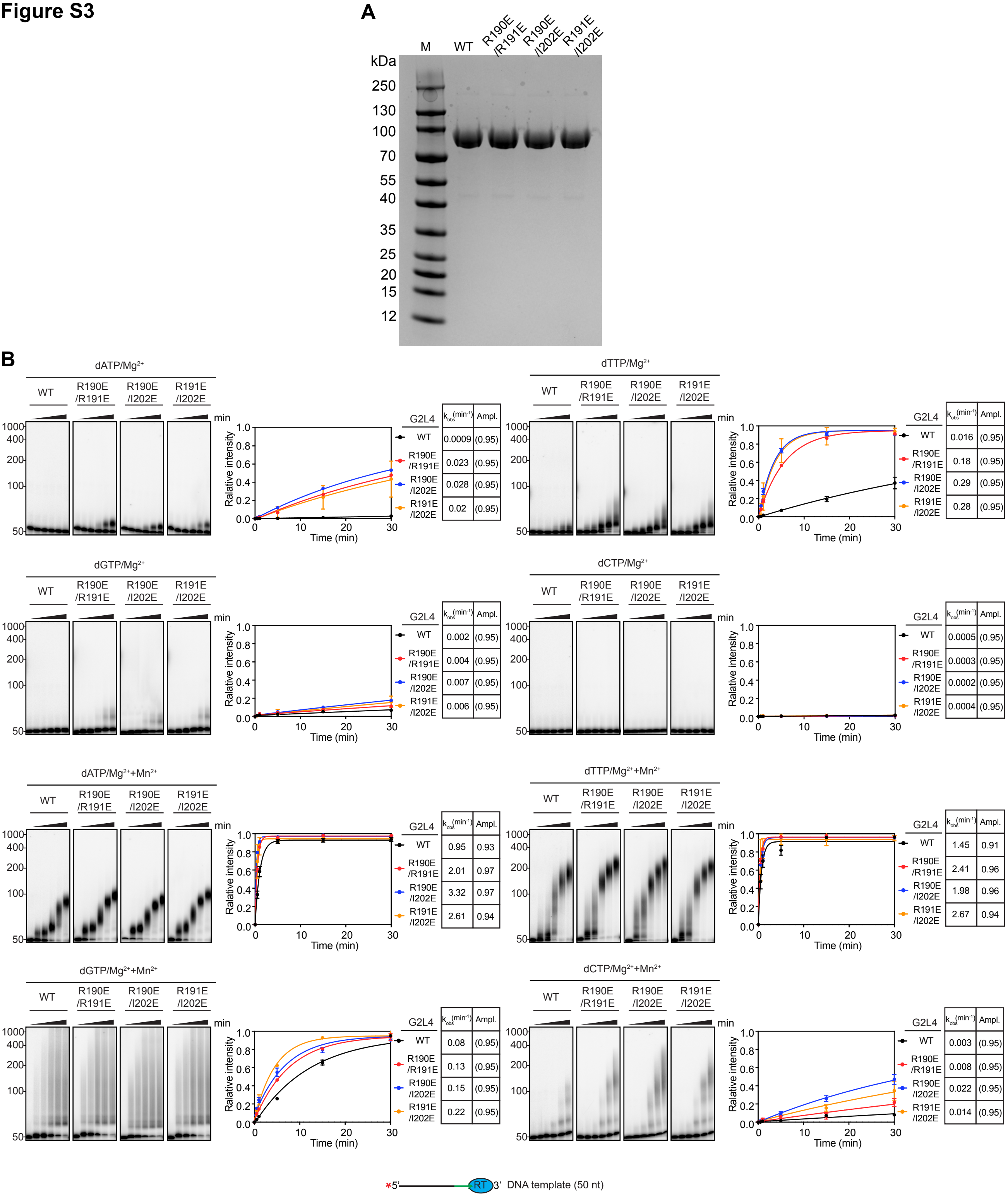
A
M
WT
R190E
/R191E
R190E
/I202E
R191E
/I202E
kDa
250
130
100
70
55
40
35
25
20
15
12
B
dATP/Mg2+
dTTP/Mg2+
dGTP/Mg2+
dCTP/Mg2+
dATP/Mg2+ +Mn2+
dTTP/Mg2+ +Mn2+
dGTP/Mg2+ +Mn2+
dCTP/Mg2+ +Mn2+
min
1000
400
200
100
50
Ralative intensity
Time (min)
G2L4
k_obs (min-1)
Ampl.
dATP/Mg2+: WT 0.0009 (0.95); R190E/R191E 0.023 (0.95); R190E/I202E 0.028 (0.95); R191E/I202E 0.02 (0.95)
dTTP/Mg2+: WT 0.016 (0.95); R190E/R191E 0.18 (0.95); R190E/I202E 0.29 (0.95); R191E/I202E 0.28 (0.95)
dGTP/Mg2+: WT 0.002 (0.95); R190E/R191E 0.004 (0.95); R190E/I202E 0.007 (0.95); R191E/I202E 0.006 (0.95)
dCTP/Mg2+: WT 0.0005 (0.95); R190E/R191E 0.0003 (0.95); R190E/I202E 0.0002 (0.95); R191E/I202E 0.0004 (0.95)
dATP/Mg2+ +Mn2+: WT 0.95 0.93; R190E/R191E 2.01 0.97; R190E/I202E 3.32 0.97; R191E/I202E 2.61 0.94
dTTP/Mg2+ +Mn2+: WT 1.45 0.91; R190E/R191E 2.41 0.96; R190E/I202E 1.98 0.96; R191E/I202E 2.67 0.94
dGTP/Mg2+ +Mn2+: WT 0.08 (0.95); R190E/R191E 0.13 (0.95); R190E/I202E 0.15 (0.95); R191E/I202E 0.22 (0.95)
dCTP/Mg2+ +Mn2+: WT 0.003 (0.95); R190E/R191E 0.008 (0.95); R190E/I202E 0.022 (0.95); R191E/I202E 0.014 (0.95)
*5'
RT
3'
DNA template (50 nt)

**Figure S3. SDS-polyacrylamide gel electrophoresis of WT and RT3a mutant G2L4 RTs and further biochemical assays related to Figure 3**

(A) Coomassie blue-stained NuPAGE 4-12% Bis-Tris gel of WT and RT3a mutant G2L4 RTs used in biochemical assay. Proteins were expressed with a N-terminal MBP tag and purified as described in Methods. The numbers to the left of the gel indicate molecular weights of a Color Prestained Broad Range (10-250 kDa) protein ladder (New England Biolabs).

(B) Terminal transferase assay time courses with a 5'-$^{32}$P-labeled 50-nt DNA substrate without a 3'-blocking group. Reactions were initiated by adding 1 mM of a single dNTP (dATP, dCTP, dGTP, and dTTP) and incubated at 37°C for times up to 30 min in reaction medium containing 10 mM $Mg^{2+}$ in the absence (top panels) or presence (bottom panels) of 1 mM $Mn^{2+}$. The numbers to the left of the gel indicate the positions of 5'-labeled RiboRuler Low Range RNA Ladder size markers run in a parallel lane. The plots to the right of the gel show the average value and variance for two repeats of the experiment. Tables to the right of the plots show the rate constants ($k_{obs}$) and amplitudes (Ampl.) of labeled products >50 nt obtained by fitting to a first-order rate equation. Ampl. values in parentheses represent fixed amplitudes for reactions that did not reach an end point based on the average Ampl. value for those that reached a clear end point during the experiment. The red asterisk in the schematic at the bottom indicates a 5'-$^{32}$P-label.

**Figure S4**

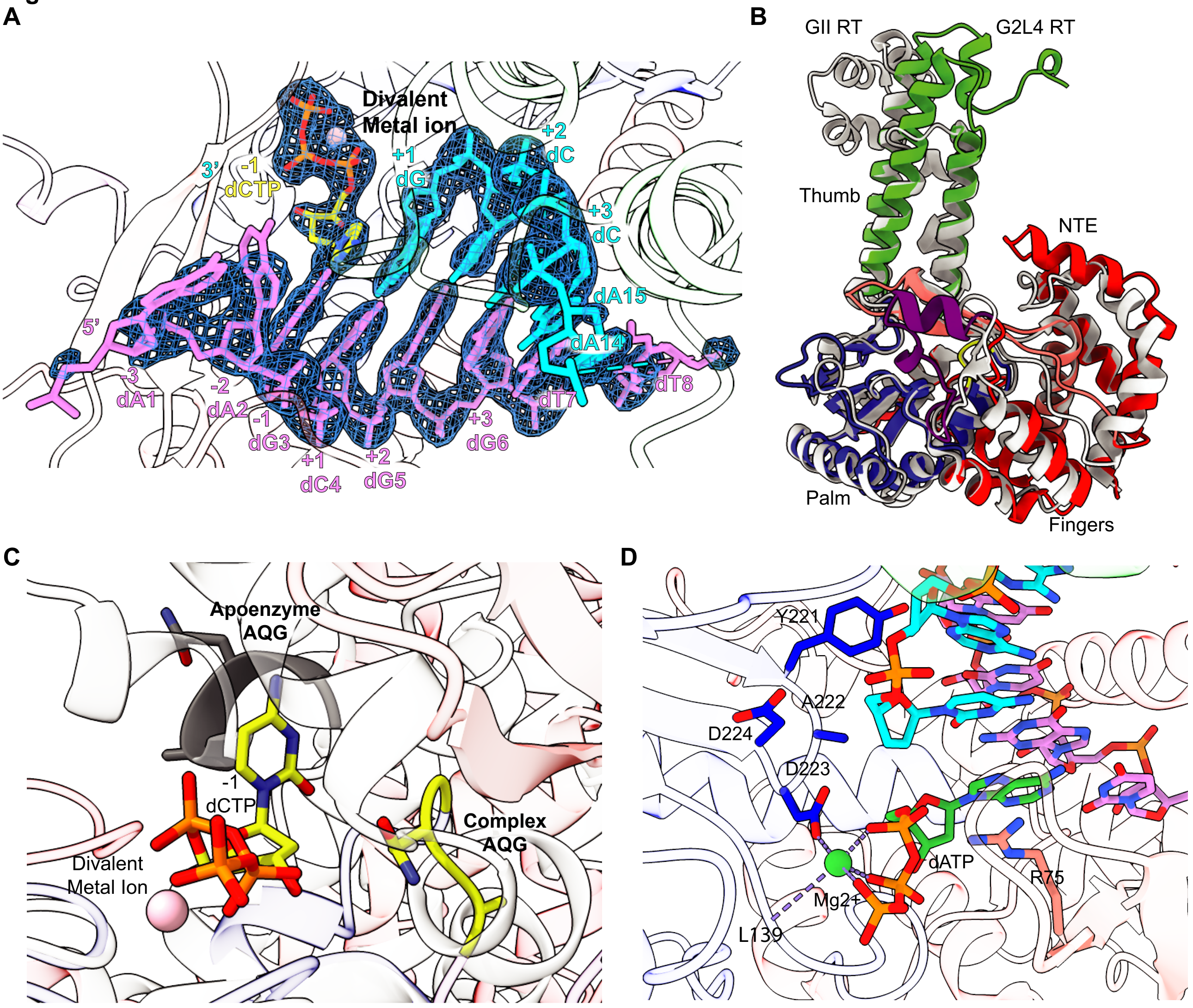
A
Divalent
Metal ion
3'
-1
dCTP
+1
dG
+2
dC
+3
dC
dA15
dA14
5'
-3
dA1
-2
dA2
-1
dG3
+1
dC4
+2
dG5
+3
dG6
dT7
dT8
B
GII RT
G2L4 RT
Thumb
NTE
Palm
Fingers
C
Apoenzyme
AQG
-1
dCTP
Divalent
Metal Ion
Complex
AQG
D
Y221
A222
D224
D223
dATP
Mg2+
R75
L139

**Figure S4. 2Fo-Fc map of G2L4 RT complex with the bound snapback DNA substate and incoming dNTP related to Figure 4**

(A) The 2Fo-Fc map was calculated by doubling the observed electron density (Fo) and subtracting the calculated electron density (Fc) to provide an estimate of the true electron density of the active site of G2L4 RT with bound substrate. Positive density of the electron cloud is shown as mesh and colored in blue contoured at a level of 1.5 sigma (σ). Nucleotides are numbered and colored as in Figure 4A.

(B) Superimposition of active conformations of G2L4 RT and GII RT. Different regions of G2L4 RT are color-coded as in Figure 1, and GII RT is colored gray.

(C) Superimposition of G2L4 RT apoenzyme and snapback DNA synthesis structure focusing on the AQG motif in the snapback DNA complex (yellow) and apoenzyme (gray). The pink sphere is a divalent metal ion in the snapback DNA complex structure.

(D) Close-up view of the active site region of GII RT[4] highlighting key residues, including D223 (blue) and L139 involved in coordination of $Mg^{2+}$ (green sphere) and binding of the incoming dATP (green base, orange triphosphates) by D223 (blue) and R75 (salmon). The DNA template (violet) and primer (cyan) are depicted as sticks.

### Figure S5

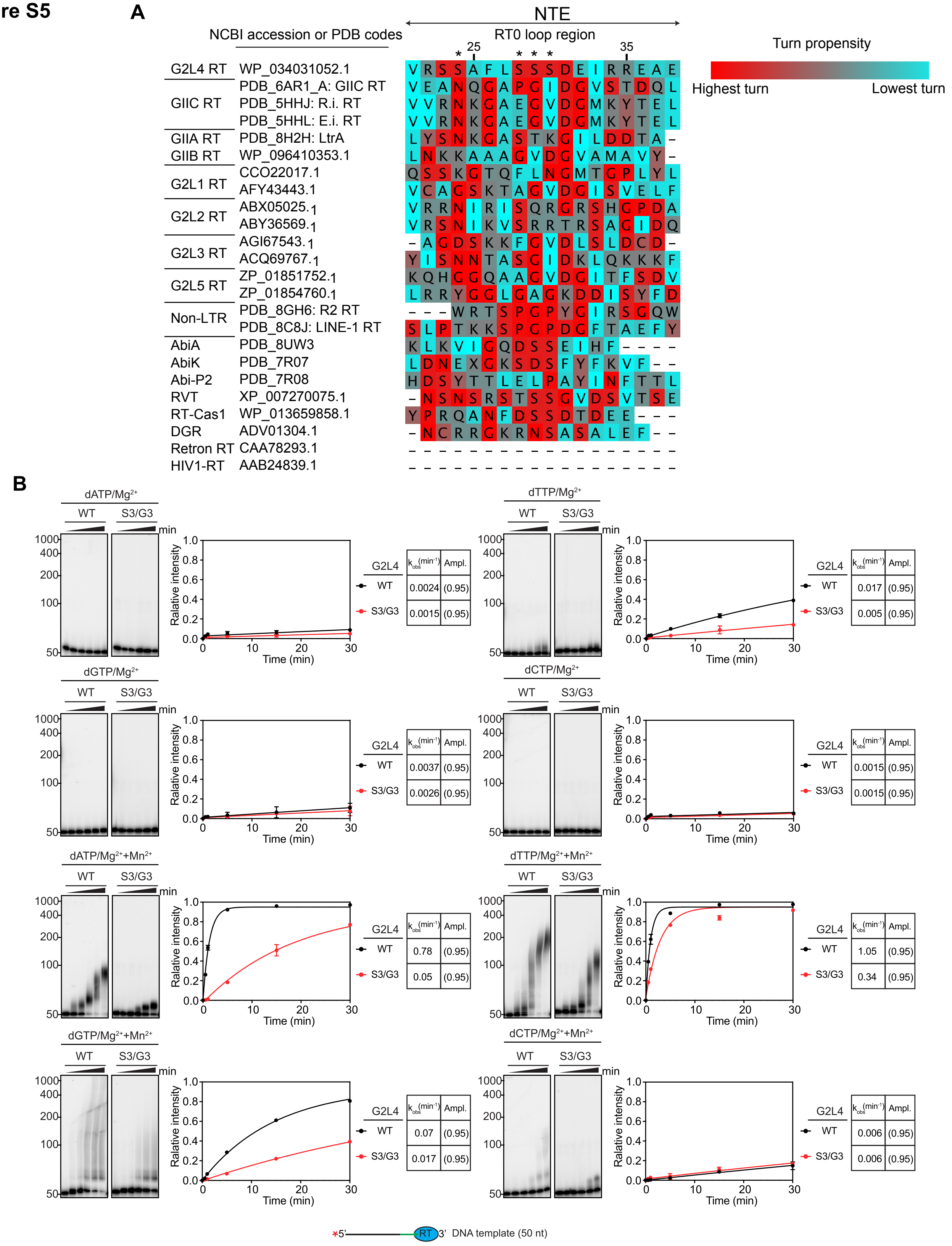
A
NTE
NCBI accession or PDB codes
RT0 loop region
25
35
Turn propensity
Highest turn
Lowest turn
G2L4 RT WP_034031052.1
GIIC RT PDB_6AR1_A: GIIC RT
PDB_5HHJ: R.i. RT
PDB_5HHL: E.i. RT
GIIA RT PDB_8H2H: LtrA
GIIB RT WP_096410353.1
G2L1 RT CCO22017.1
AFY43443.1
G2L2 RT ABX05025.1
ABY36569.1
G2L3 RT AGI67543.1
ACQ69767.1
G2L5 RT ZP_01851752.1
ZP_01854760.1
Non-LTR PDB_8GH6: R2 RT
PDB_8C8J: LINE-1 RT
AbiA PDB_8UW3
AbiK PDB_7R07
Abi-P2 PDB_7R08
RVT XP_007270075.1
RT-Cas1 WP_013659858.1
DGR ADV01304.1
Retron RT CAA78293.1
HIV1-RT AAB24839.1
B
dATP/Mg2+
dTTP/Mg2+
dGTP/Mg2+
dCTP/Mg2+
dATP/Mg2+ +Mn2+
dTTP/Mg2+ +Mn2+
dGTP/Mg2+ +Mn2+
dCTP/Mg2+ +Mn2+
WT
S3/G3
min
1000
400
200
100
50
Ralative intensity
Time (min)
G2L4
kobs (min-1)
Ampl.
dATP/Mg2+: WT 0.0024 (0.95); S3/G3 0.0015 (0.95)
dTTP/Mg2+: WT 0.017 (0.95); S3/G3 0.005 (0.95)
dGTP/Mg2+: WT 0.0037 (0.95); S3/G3 0.0026 (0.95)
dCTP/Mg2+: WT 0.0015 (0.95); S3/G3 0.0015 (0.95)
dATP/Mg2+ +Mn2+: WT 0.78 (0.95); S3/G3 0.05 (0.95)
dTTP/Mg2+ +Mn2+: WT 1.05 (0.95); S3/G3 0.34 (0.95)
dGTP/Mg2+ +Mn2+: WT 0.07 (0.95); S3/G3 0.017 (0.95)
dCTP/Mg2+ +Mn2+: WT 0.006 (0.95); S3/G3 0.006 (0.95)
*5' RT 3' DNA template (50 nt)

**Figure S5. Sequence alignments of the NTE/RT0 loop of G2L4 RT compared to other RTs and terminal transferase assays of WT and RT0 loop mutant G2L4 RTs related to Figure 5**

(A) Multiple sequence alignments of the NTE/RT0 loop region of G2L4 RT (top) with those of other RTs (NCBI accession numbers or PDB codes to the right of protein names) performed by T-Coffee via Jalview[48,50]. The color scheme of amino acids boxes is based on their Chou-Fasman two turn propensity (red, highest turn propensity; cyan, lowest turn propensity)[65]. The numbers at the top indicate the positions of the G2L4 RT sequence. The asterisks indicate conserved serine residues in G2L4 RTs.

(B) Terminal transferase assays of MBP-tagged G2L4 wild-type and S3/G3 mutant RTs in the presence of 10 mM $Mg^{2+}$ (left assays) or 10 mM $Mg^{+2}$ + 1 mM $Mn^{2+}$ (right assays) using a 5'-labeled 50-nt DNA substrate, as described in Figures S3B. Reactions were initiated by adding 1 mM of a single dNTP (dATP, dCTP, dGTP, and dTTP) and incubated at 37°C for times up to 30 min. The numbers to the left of the gels indicate the positions of 5'-labeled RiboRuler Low Range RNA Ladder size markers run in a parallel lane. The plots show the average values and variance for two repeats of the experiment. The Tables to the right of the plots indicate rate constants ($k_{obs}$) and amplitudes (Ampl.) for synthesized DNA products by WT and S3/G3 RT0 loop mutant G2L4 RTs with a curve fit to a first-order rate equation. Ampl. values in parentheses represent fixed amplitudes for reactions that did not reach an end point based on the average Ampl. value for those that reached a clear end point during the experiment. The red asterisk in the schematic at the bottom indicates a 5'-$^{32}$P-label.

### Figure S6

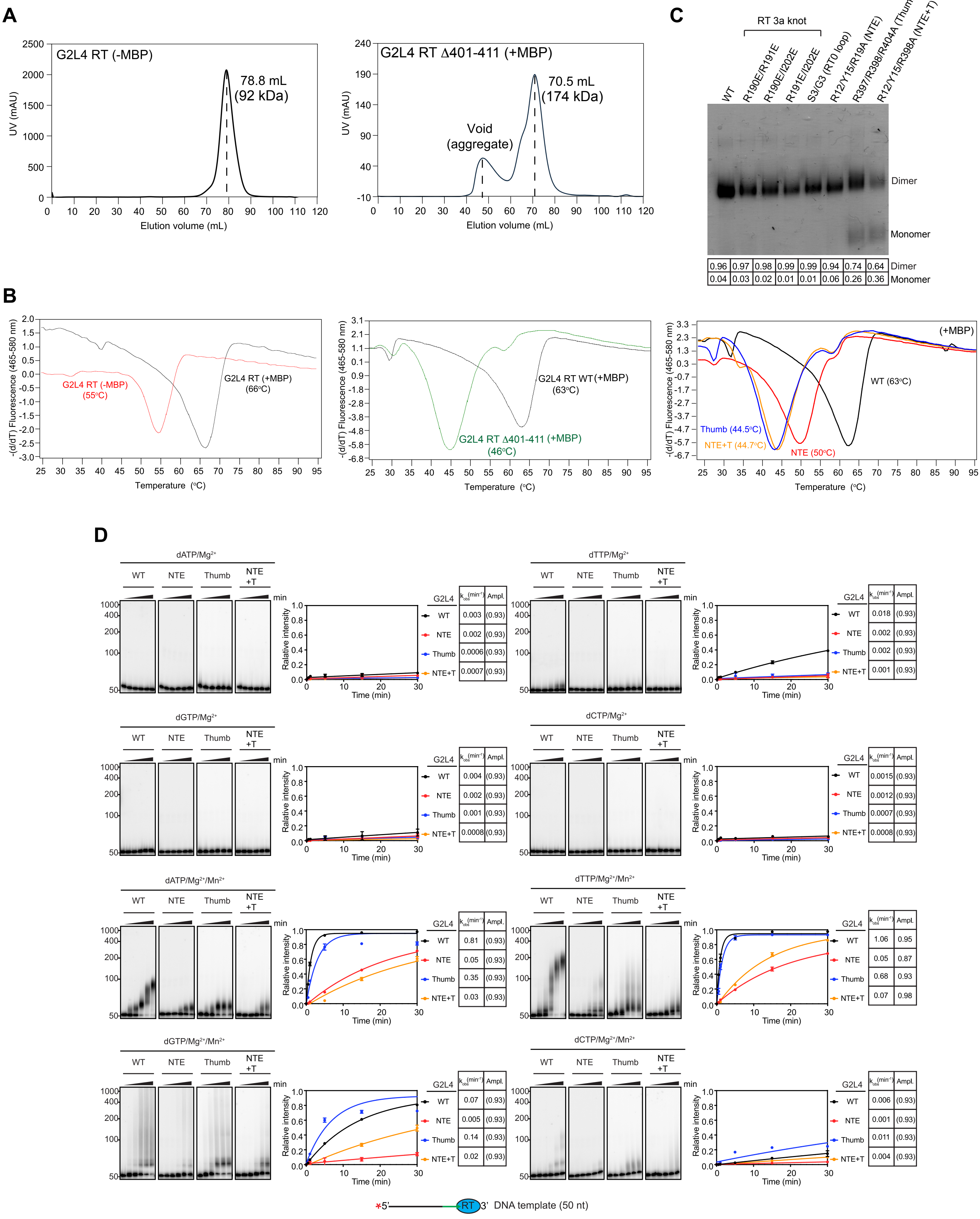

A
G2L4 RT (-MBP)
78.8 mL
(92 kDa)
UV (mAU)
Elution volume (mL)
G2L4 RT Δ401-411 (+MBP)
70.5 mL
(174 kDa)
Void
(aggregate)
C
RT 3a knot
WT
R190E/R191E
R190E/I202E
R191E/I202E
S3/G3 (RT0 loop)
R12/Y15/R19A (NTE)
R397/R398/R404A (Thumb)
R12/Y15/R398A (NTE+T)
Dimer
Monomer
0.96 0.97 0.98 0.99 0.99 0.94 0.74 0.64 Dimer
0.04 0.03 0.02 0.01 0.01 0.06 0.26 0.36 Monomer
B
-(d/dT) Fluorescence (465-580 nm)
Temperature (°C)
G2L4 RT (-MBP)
(55°C)
G2L4 RT (+MBP)
(66°C)
G2L4 RT WT(+MBP)
(63°C)
G2L4 RT Δ401-411 (+MBP)
(46°C)
(+MBP)
WT (63°C)
Thumb (44.5°C)
NTE+T (44.7°C)
NTE (50°C)
D
dATP/Mg2+
dTTP/Mg2+
dGTP/Mg2+
dCTP/Mg2+
dATP/Mg2+/Mn2+
dTTP/Mg2+/Mn2+
dGTP/Mg2+/Mn2+
dCTP/Mg2+/Mn2+
WT
NTE
Thumb
NTE +T
min
Relative intensity
Time (min)
G2L4 kobs(min-1) Ampl.
dATP/Mg2+: WT 0.003 (0.93); NTE 0.002 (0.93); Thumb 0.0006 (0.93); NTE+T 0.0007 (0.93)
dTTP/Mg2+: WT 0.018 (0.93); NTE 0.002 (0.93); Thumb 0.002 (0.93); NTE+T 0.001 (0.93)
dGTP/Mg2+: WT 0.004 (0.93); NTE 0.002 (0.93); Thumb 0.001 (0.93); NTE+T 0.0008 (0.93)
dCTP/Mg2+: WT 0.0015 (0.93); NTE 0.0012 (0.93); Thumb 0.0007 (0.93); NTE+T 0.0008 (0.93)
dATP/Mg2+/Mn2+: WT 0.81 (0.93); NTE 0.05 (0.93); Thumb 0.35 (0.93); NTE+T 0.03 (0.93)
dTTP/Mg2+/Mn2+: WT 1.06 0.95; NTE 0.05 0.87; Thumb 0.68 0.93; NTE+T 0.07 0.98
dGTP/Mg2+/Mn2+: WT 0.07 (0.93); NTE 0.005 (0.93); Thumb 0.14 (0.93); NTE+T 0.02 (0.93)
dCTP/Mg2+/Mn2+: WT 0.006 (0.93); NTE 0.001 (0.93); Thumb 0.011 (0.93); NTE+T 0.004 (0.93)
*5' RT 3' DNA template (50 nt)

**Figure S6. Profiling of G2L4 RT dimer interface mutants related to Figure 6**

(A) Size-exclusion chromatography of WT G2L4 RTs with a cleaved MBP tag (left panel) and a G2L4 RT C-terminal deletion mutant (Δ401-411 amino acids; right panel). The molecular weights in parenthesis were calculated based on the elution volume of the peak (dashed line) relative to the protein standard calibration graph of Figure S1A.

(B) Differential Scanning Fluorimetry (DSF) assays. Left panel, G2L4 RT with (black) or without (red) an MBP tag; middle panel, C-terminal deletion mutant of G2L4 RT (Δ401-411 amino acids, green) versus WT G2L4 RT (black); right panel, dimer interface NTE mutant (R12/Y15/R19A; red); Thumb (T) domain mutant (R397/R398/R404A; blue); and NTE+T mutant (R12/Y15/R398A; yellow). The plots show the derivative of fluorescence intensity as a function of temperature, highlighting transitions or melting points indicative of structural changes or stability shifts.

(C) Coomassie blue-stained non-denaturing gel for WT and mutant G2L4 RTs with an N-terminal MBP tag. Dimer and monomer bands are labeled to the right. The fractions of monomer bands indicated below the gel were calculated by using ImageQuant TL 10.2 software.

(D) Terminal transferase assays with or without $Mn^{2+}$ of MBP-tagged G2L4 RT dimer interface mutants using a 5'-labeled 50-nt DNA substrate, as described in Figure S3B.

**Figure S7**

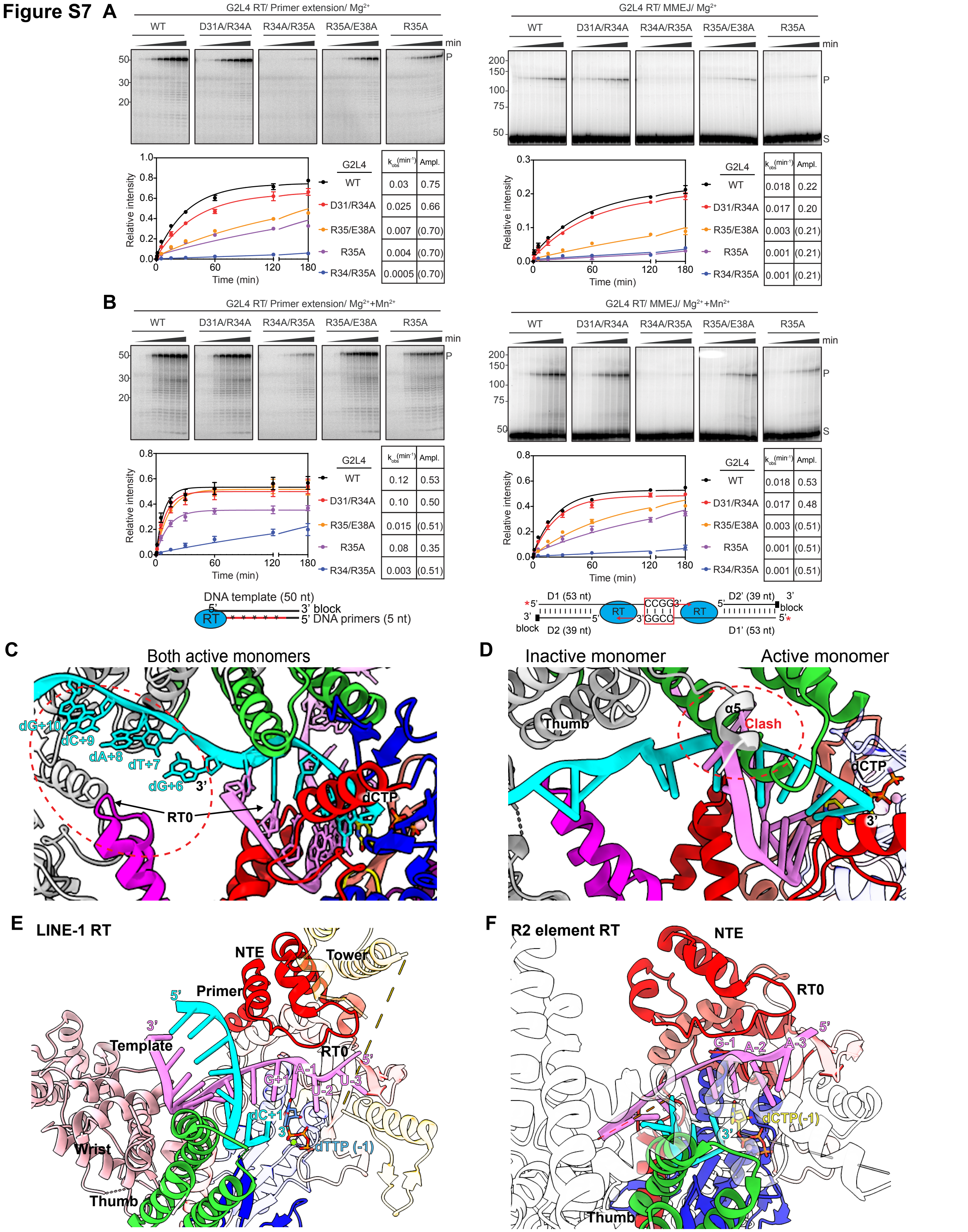

A
G2L4 RT/ Primer extension/ Mg2+
WT
D31A/R34A
R34A/R35A
R35A/E38A
R35A
min
50
30
20
P
Relative intensity
Time (min)
G2L4
kobs(min-1)
Ampl.
WT 0.03 0.75
D31/R34A 0.025 0.66
R35/E38A 0.007 (0.70)
R35A 0.004 (0.70)
R34/R35A 0.0005 (0.70)
G2L4 RT/ MMEJ/ Mg2+
200
150
100
75
50
S
G2L4
kobs(min-1)
Ampl.
WT 0.018 0.22
D31/R34A 0.017 0.20
R35/E38A 0.003 (0.21)
R35A 0.001 (0.21)
R34/R35A 0.001 (0.21)
B
G2L4 RT/ Primer extension/ Mg2++Mn2+
G2L4
kobs(min-1)
Ampl.
WT 0.12 0.53
D31/R34A 0.10 0.50
R35/E38A 0.015 (0.51)
R35A 0.08 0.35
R34/R35A 0.003 (0.51)
G2L4 RT/ MMEJ/ Mg2++Mn2+
G2L4
kobs(min-1)
Ampl.
WT 0.018 0.53
D31/R34A 0.017 0.48
R35/E38A 0.003 (0.51)
R35A 0.001 (0.51)
R34/R35A 0.001 (0.51)
DNA template (50 nt)
5'
3' block
RT
5' DNA primers (5 nt)
D1 (53 nt)
D2' (39 nt)
D2 (39 nt)
D1' (53 nt)
CCGG
GGCC
block
C
Both active monomers
dG+10
dC+9
dA+8
dT+7
dG+6
3'
RT0
dCTP
D
Inactive monomer
Active monomer
Thumb
α5
Clash
dCTP
3'
E
LINE-1 RT
NTE
Tower
Primer
5'
3'
Template
RT0
G+1
A-1
U-2
U-3
dC+1
dTTP (-1)
Wrist
Thumb
F
R2 element RT
NTE
RT0
G-1
A-2
A-3
dCTP(-1)
3'
Thumb

**Figure S7. G2L4 RT biochemical assay and models supporting the suggested G2L4 RT MMEJ mechanism and RT0 loop interactions for human LINE-1 and insect R2 non-LTR-retroelements related to Figure 7.**

(A and B) Biochemical assays of G2L4 RT NTE mutants in reaction media containing 10 mM $Mg^{2+}$ (panel A) or 10 mM $Mg^{2+}$ plus 1 mM $Mn^{2+}$ (panel B). Left, primer extension assays; right, MMEJ assays. The plots show the average values and variance for two repeats of the experiment. Ampl. values in parentheses represent fixed amplitudes for reactions that did not reach an end point based on the average Ampl. value for those that reached a clear end point during the experiment.

(C) Model of a G2L4 RT dimer with two active monomers bound to a 5-bp annealed microhomology between the single-stranded 3' overhangs from the left side (cyan) and right side (violet) of the DSB. A dashed red circle highlights a gap between single-stranded region of the MMEJ substrate and the trailing active monomer.

(D) Model of a G2L4 RT dimer with an active and inactive monomers bound to a 7-bp annealed microhomology between the same single-strand 3' overhangs as panel C. A steric clash is highlighted in a dashed red circle.

(E) Wide-angle views of the RT0 region of LINE-1 RT (PDB: 8C8J)[15] with bound RNA template/DNA primer substrate and dTTP. The RNA template (violet), DNA primer (cyan), and bound dTTP (light blue/orange) are depicted as sticks. Tower (light yellow) and Wrist (pink) are additional regions of LINE-1 RT.

(F) Wide-angle views of the RT0 region of *Bombyx mori* R2 element RT (PDB:8GH6)[38] modeled with the same DNA temple/DNA primer substrate as G2L4 RT. The DNA template (violet), DNA primer (cyan), and bound dTTP (blue base) and dCTP (yellow base) are depicted as sticks.

### Table S1. Oligonucleotides

| Use | Name | Sequence |
|---|---|---|
| Cloning | G2L4 RT R190E Top | 5' AACATTGATGACCTTGAACGCAAGGGACTGGATGAA 3' |
| | G2L4 RT R191E Top | 5' AACATTGATGACCTTCGCGAAAAGGGACTGGATGAA 3' |
| | G2L4 RT R190E/R191E Top | 5’ AACATTGATGACCTTGAAGAAAAGGGACTGGATGAA 3’ |
| | G2L4 RT R190E/R191E Bot | 5’ CGCCAGATCCGTGCGGATTGCCTCTTCAAAAAGCTT 3’ |
| | G2L4 RT I202E Top | 5' GAAATCTTTCCCATTGGCGAAGAAGGGGTTGCTCAA 3' |
| | G2L4 RT I202E Bot | 5' ATCCAGTCCCTTGCGGCGAAGGTCATCAATGTTCGC 3' |
| | G2L4 RT S3/G3 Top | 5' TCGGCGTTTTTGGGTGGTGGTGATGAGATCCGT 3' |
| | G2L4 RT S3/G3 Bot | 5' GGAGCGAACCTTACGCCATGCACCGTACAGGGT 3' |
| | G2L4 RT R19A Top | 5' TCCTCGGCGTTTTTGTCTTCCAGTGATGAGATCCGTCGTGAA 3' |
| | G2L4 RT R19A Bot | 5' GCGAACCTTTTCCCATGCACCTGCCAGGGTTTCCAGGGAGCA 3' |
| | G2L4 RT R397/398/404A (Thumb) Top | 5' ACAGTAGCAGCACGTGTATTAGGCGTGGCATTGCTTATT 3' |
| | G2L4 RT R397/398/404A (Thumb) Bot | 5' ACTGTCGCTGTTGGCAAGAATGGCAATTTTTTC 3' |
| | G2L4 RT R12/Y15A Top | 5' CGTAAGGTTCGCTCCTCGGCGTTTTTGTCTTCC 3' |
| | G2L4 RT R12/Y15A Bot | 5' CCATGCACCCGCCAGGGTCGCCAGGGAGCAAAT 3' |
| | G2L4 RT R398A Top | 5' GACAGTACAGTACGTGCCCGTGTATTAGGCGTG 3' |
| | G2L4 RT R398A Bot | 5' GCTGTTGGCAAGAATGGCAATTTTTTCCGCTTC 3' |
| | G2L4 RT C-term deletion (401-411) Top | 5' TGAGGATCCGAATTCCCTGCAGGTA 3' |
| | G2L4 RT C-term deletion (401-411) Bot | 5' TACACGACGACGTACTGTACTGTCGC 3' |
| | G2L4 RT D31A/R34A Top | 5’ ATCGCGCGTGAAGCAGAGGAATTTGAGAG 3’ |
| | G2L4 RT D31A/R34A Bot | 5’ CTCCGCACTGGAAGACAAAAACGCCGAG 3’ |
| | G2L4 RT R34A/R35A Top | 5’ TGATGAGATCGCAGCAGAAGCAGAGGAATTTGAGAGTCGTTTGC 3’ |
| | G2L4 RT R34A/R35A Bot | 5’ CTGGAAGACAAAAACGCCGAGG 3’ |
| | G2L4 RT R35A/E38A Top | 5’ GCAGCAGAATTTGAGAGTCGTTTGCCCGAC 3’ |
| | G2L4 RT R35A/E38A Bot | 5’ TTCCGCACGGATCTCATCACTGGAAGAC 3’ |
| | G2L4 RT R35A Top | 5’ TGAGATCCGTGCGGAAGCAGAGGAATTTG 3’ |
| | G2L4 RT R35A Bot | 5’ TCACTGGAAGACAAAAACGCCG 3’ |
| Snapback/ Terminal transferase assays | 50-nt DNA | 5' GCAATAATCTATACAATACAACACATACAAACAAATTCTTAAGGTCCCAA 3' |
| Primer extension assays | 50-nt DNA 3' block (Inverted dT at the 3'-end) | 5' GCAATAATCTATACAATACAACACATACAAACAAATTCTTAAGGTCCCAA/InvdT/ 3' |
| | 5-nt DNA primer | 5' TTGGG 3' |
| MMEJ assays | 53-nt DNA 4 microhomology (CCGG) Top | 5' CCCTGTACAGTAAGAGCCTACTCATGGATCCTCCTTGTGATGTAAGGTCCCGG 3' |
| | 39-nt DNA 3' block (Inverted dT at the 3'-end) Bot | 5' ACAAGGAGGATCCATGAGTAGGCTCTTACTGTACAGGG/ InvdT / 3' |
| Snapback substrate for X-ray crystallography | 15-nt snapback DNA | 5’ AAGCGGTTAACCCAA 3’ |

**Table S2. Recombinant plasmids**

| Name | Source |
|---|---|
| pMal-G2L4 RT WT | Park et al., 2022[20] |
| pMal-G2L4 RT R190E/R191E | This Study |
| pMal-G2L4 RT R190E/I202E | This Study |
| pMal-G2L4 RT R191E/I202E | This Study |
| pMal-G2L4 RT S3/G3 | This Study |
| pMal-G2L4 RT R12/Y15/R19A (NTE) | This Study |
| pMal-G2L4 RT R397/R398/R404A (Thumb) | This Study |
| pMal-G2L4 RT R12/Y15/R398A (NTE+T) | This Study |
| pMal-G2L4 RT D31A/D34A | This Study |
| pMal-G2L4 RT D34A/D35A | This Study |
| pMal-G2L4 RT D35A/D38A | This Study |
| pMal-G2L4 RT D35A | This Study |
